# Effect of Voluntary Adolescent Alcohol Consumption on Encoding of Decision-Related Variables in Prefrontal Cortex

**DOI:** 10.1101/2021.03.09.434599

**Authors:** Samantha Corwin, Jamie D Roitman

## Abstract

The medial and orbitofrontal regions of prefrontal cortex (PFC) have been implicated in guiding optimal behavior and updating the economic value of rewards that result from choice behaviors. Both regions mature through adolescence into early adulthood and are thus vulnerable to exposure to neurotoxins, such as alcohol, during this critical developmental window. We sought to examine how voluntary alcohol consumption during adolescence would alter long-term PFC function and subsequent decision-making behavior in adulthood. Male and female rats were given adolescent intermittent ethanol (AIE) exposure to provide voluntary access to alcohol during the period of PFC maturation. In adulthood, we assessed the long-term effects on decision-making behavior using a risk task in adulthood, while concurrently recording neural activity in orbitofrontal cortex (OFC) and medial prefrontal cortex (mPFC). While control animals’ preferences for risky rewards increased with the likelihood of their delivery, AIE animals showed an overall reduction in their preferences for the risky option with higher levels of alcohol consumption, suggesting reduced discriminability of uncertain rewards and a shift away from the potential for reward omission. During task performance, neurons in mPFC and OFC responded to events (lever press, reward delivery). In mPFC, neurons with phasic increases at the time of lever press showed a reversal from larger elevations for risky presses in control animals to larger elevations for certain presses as prior alcohol consumption increased. Neurons in mPFC generally showed less discrimination of reward outcome with increased alcohol consumption as well. In OFC, responses to lever press were largely unaffected by AIE exposure. However, encoding of reward size in OFC showed differential effects in males and females. With higher alcohol consumption in males, OFC neurons showed largest excursions from baseline activity in response to largest reward, and smallest excursions for reward omission. This discrimination was reduced as prior alcohol consumption increased. In females, neurons with increased reward-activity, showed an overall higher level of activity due to stronger responses to certain rewards that were selected more frequently. Collectively the results show diminished capacity of PFC to encode decision-related elements to guide adaptive behavior and further clarify the lasting impact of adolescent alcohol use on neural function and behavior, even in the absence of continued use.

## INTRODUCTION

Prefrontal cortex (PFC) has been long implicated in directing decisions, particularly when choices must take uncertainty into account. Neural activity in subregions of PFC, such as medial prefrontal cortex (mPFC) and orbitofrontal cortex (OFC), have been shown to be necessary for advantageous decision-making, including reward evaluation and cue sensitivity (Orsini et al., 2015). Due to the plasticity of PFC maturation in adolescents, they are more susceptible to anatomical and functional disruption of this area due to neurotoxin exposure, resulting in permanent damage. However, it is not understood how chronic insult to PFC during development may differentially affect males and females, as well as different sub-regions of PFC.

Alcohol is the most widely used intoxicating substance among youths and initial experimentation with alcohol often transpires during ongoing maturation of PFC. Adolescents frequently engage in risky behaviors, such as binge drinking, that result in immediate and lasting consequences. Immediate impacts range from reduced verbal and spatial memory (Brown et al., 2000) to impaired decision-making (McMurray et al., 2014). Previous studies have demonstrated long-term repercussions of adolescent drinking on male rats’ risk preference and OFC function (Boutros et al., 2015; Clark et al., 2012; McMurray et al., 2015; N. a Nasrallah, Yang, & Bernstein, 2009) however, whether these results are similar in females has yet to be determined. Furthermore, it is also not known how adolescent alcohol use might differentially affect subregions of PFC. Additionally, adolescence typically marks the onset of addiction and other psychiatric disorders, providing further incentive to evaluate the long-term effects of adolescent alcohol use on neural activity.

We tested how different sub regions of PFC encode decision-related variables and how decisions that are dependent on PFC activity are altered by variable alcohol consumption in males and females. Animals were given adolescent intermittent ethanol (AIE) exposure to a sweetened gelatin containing 10% alcohol or no alcohol. After prolonged abstinence, rats were trained and tested in adulthood on a risk task in which they had the choice between small-certain rewards or large-risky rewards. The rats’ sensitivity to the changing frequency of the large reward’s delivery and their preference for the larger, riskier reward was measured by exposing them to an array of reward contingencies. While they performed the behavioral task, we recorded from individual neurons in mPFC and OFC to examine changes in decision-related activity.

Extracellular, single cell, electrophysiological recordings were used to record action potentials in OFC and mPFC and align them to events that occurred during the risk task. Thus, based on the timing and patterns of activity, it could be established what specific role these regions were playing in the decision. Additionally, by measuring discrepancies in neural activity in rats who had consumed alcohol in adolescence compared to those who had not, it could be ascertained how alcohol was changing the neural basis of behavior.

We hypothesized that both OFC and mPFC would both contain neurons that altered their firing rates at the time of the lever press or at the time of reward delivery. However, we expected mPFC to be more active at the time of lever press and OFC to show greater modulation during reward encoding, given their proposed roles in direction of behavior and reward evaluation. We hypothesized increased preference for the risky option in animals that consumed more alcohol would correspond with altered task-related responses in PFC. Neurons that responded during reward delivery were expected to exhibit alcohol-dependent hypoactivity regardless of sex. It was predicted that this blunted activity would result in the inability to discriminate different reward sizes through firing rate. Previous research has shown stunted OFC activity in high alcohol consuming male rats (McMurray et al., 2015) and a similar result was anticipated in females. We expected responses at the time of lever press to reflect the animals’ changing preferences for the behavioral options available.

## MATERIALS AND METHODS

### Animals

Male and female Long-Evans rats were acquired from Charles River Laboratory (Wilmington, MA) on PD (postnatal day) 22. Rats were housed in same-sex groups of 3 in polycarbonate cages (45×24×20 cm) and were provided with ad libitum chow (LabDiet 5012 Richmond, IN) and water, unless otherwise noted. The colony was maintained on a 12:12 light/dark cycle and electrophysiological recordings were conducted during the light cycle. Animals were treated under the approval of the Animal Care Committee of the University of Illinois at Chicago and in accordance with the National Institutes of Health’s guidelines.

### Behavioral Methods

#### Adolescent Intermittent Ethanol (AIE) exposure

The timeline of the overall study is shown in Figure 1. From PD 30 to PD 51. Rats were given access to alcohol in the form of a “gelatin shot” that was comprised of 2.5% Knox gelatin (Kraft Foods, Northfield, IL), 10% Polycose (SolCarb, Pawcatuck, CT), and 10% EtOH (190 proof) by weight. Control animals were given non-alcoholic gelatin that contained additional polycose to match the caloric content of alcohol in the experimental gelatin. In each group of 3 animals, 2 received alcohol in the gelatin and the third received control gelatin. Rats had access to gelatin using a modified ‘drinking in the dark’ paradigm (Bell et al., 2011; McMurray et al., 2015) that allowed them to consume the gelatin at the beginning of the dark cycle. Rats were given intermittent access to the gelatin throughout adolescence (2 days on, 2 days off, 2 days on, 1 day off) starting with 24 hours of access on PD 30 and PD 31, 6 hours on PD 34, 3 hours on PD 35, and 2 hours on all subsequent access days until PD 51. Before the gelatin became available for consumption, rats were weighed and transferred from their home cage to a larger polycarbonate cage (52 x 28 x 21 cm) that was equipped with mesh dividers to ensure that rats could only access to their own gelatin while still maintaining social contact with cage mates. Rats had ongoing access to chow and water while housed in the larger apparatus. Jars of prepared gelatin were weighed before and after the access periods to determine the amount consumed.

**Figure 1.**
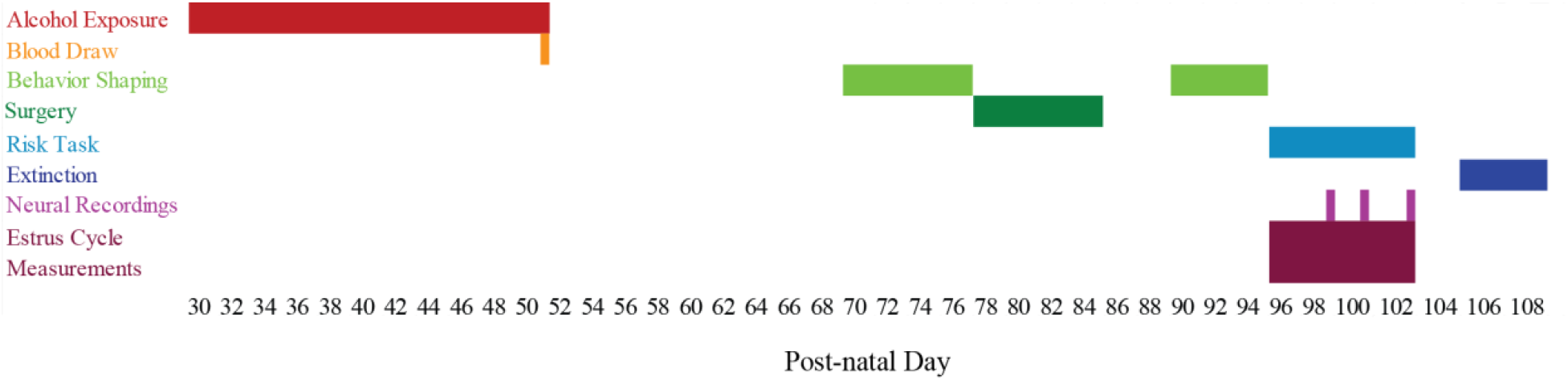
Timeline to assess the effect of adolescent alcohol use on decision-behavior and neural activity. Rats had intermittent access to alcoholic or control gelatin from PD 30-51, tail blood was drawn on PD51 to assess BEL. On PD 70 rats were trained to lever press for sugar pellets until they reached criteria. Upon completion, around PD 75, rats had microelectrodes surgically implanted in OFC and mPFC. They were allowed at least one week to recover from surgery. Around PD 90 rats were reintroduced to the task in the form of magnitude discrimination for a maximum of 5 days. Immediately upon completion of magnitude discrimination, rats were tested on the risk task and subsequent extinction sessions. Estrous cycle stage measurements were taken daily during days that they were performing the risk task.

#### Blood ethanol levels

Blood was collected via tail nick on the last day of AIE (Figure 1; PD 51) and emulsified with heparin. Blood plasma was extracted from each sample and blood ethanol levels (BEL) were measured at a later time using an AM1 alcohol analyzer (Analox).

#### Behavioral training and risk task

Behavior training and testing were conducted in operant chambers (35 x 32 x 33 cm) equipped with a central pellet dispenser, two levers on either side of the pellet dispenser, one cue light above each lever, a house light in the back of the chamber, and an overhead fan (Med Associates St Albans, VT). An infrared beam was positioned at the opening of the pellet dispenser to mark head entries into the area. Rats were food restricted to no less than 90% of free-feeding bodyweight during behavior shaping and the risk task.

##### Magazine Training

On PD 70 rats began magazine training (Figure 1), which consisted of the dispensing of sugar pellets (45mg; BioServ, Frenchtown, NJ) at variable intervals (20, 30, 60, or 90 s) for 45 minutes. Magazine training continued daily until rats ate 90% of the dispensed sugar pellets for 2 consecutive days. Rats were then trained to lever press for sugar pellets using a fixed ratio of 1 (FR-1) schedule containing a forced choice block (12 trials per lever) and a free choice block (88 trials). The FR-1 schedule continued until rats completed 50% of trials for 2 days. Side bias during FR-1 training was determined to be present if a rat pressed one lever for more than 70% of completed trials. If a side bias was observed, the favored lever would be designated as the ‘small, certain’ lever during magnitude discrimination and the risk task. This ensured that any initial spatial preference was superseded by the preference for the large reward. If a bias was not observed, then the small, certain lever was assigned at random.

##### Magnitude Discrimination

Magnitude discrimination consisted of a block of forced-choice trials (12 trials) and a block of free-choice trials (88 trials), wherein one lever delivered a large payout (three 45mg sucrose pellets) when it was pressed (varied ratio of 1: VR-1) and the other lever delivered a small payout (one 45mg sucrose pellet) when it was pressed (fixed ratio of 1: FR-1). Rats performed magnitude discrimination until they selected the VR-1 lever on 70% of completed free-choice trials for 2 consecutive days or for a maximum of 5 consecutive days.

##### Risk Task

Once rats successfully completed magnitude discrimination, they then performed two days of an “anchor session” for the risk task in which the VR-1 lever’s ratio of delivery changed to 3 (VR-3). This transformed it to the ‘large, risky’ lever and when pressed, it resulted in the delivery of 3 sugar pellets 33% of the time (VR-3); the intention of this was to expose rats to the uncertainty aspect of the task without contaminating the data with carry-over effects caused by the risky lever bias evoked during magnitude discrimination.

During the risk task, the lever associated with the FR-1 schedule during magnitude discrimination was designated as the ‘small, certain’ payoff lever (one 45mg sucrose pellet, 100% of the time) and the VR-1 lever became associated with a potentially large, risky reward (zero or three 45mg sucrose pellets, 16%, 33%, or 66% of the time). Each session began with a block of 12 forced trials per lever followed by 76 free choice trials. The rats were tested on each probability for 2 consecutive days but only data from the second day were analyzed to exclude trials in which the animals were first experiencing the new probability. The large, risky lever probabilities were ordered pseudo-randomly to ensure the 33% session did not immediately follow the two days of “anchor sessions” (Figure 2A).

**Figure 2.**
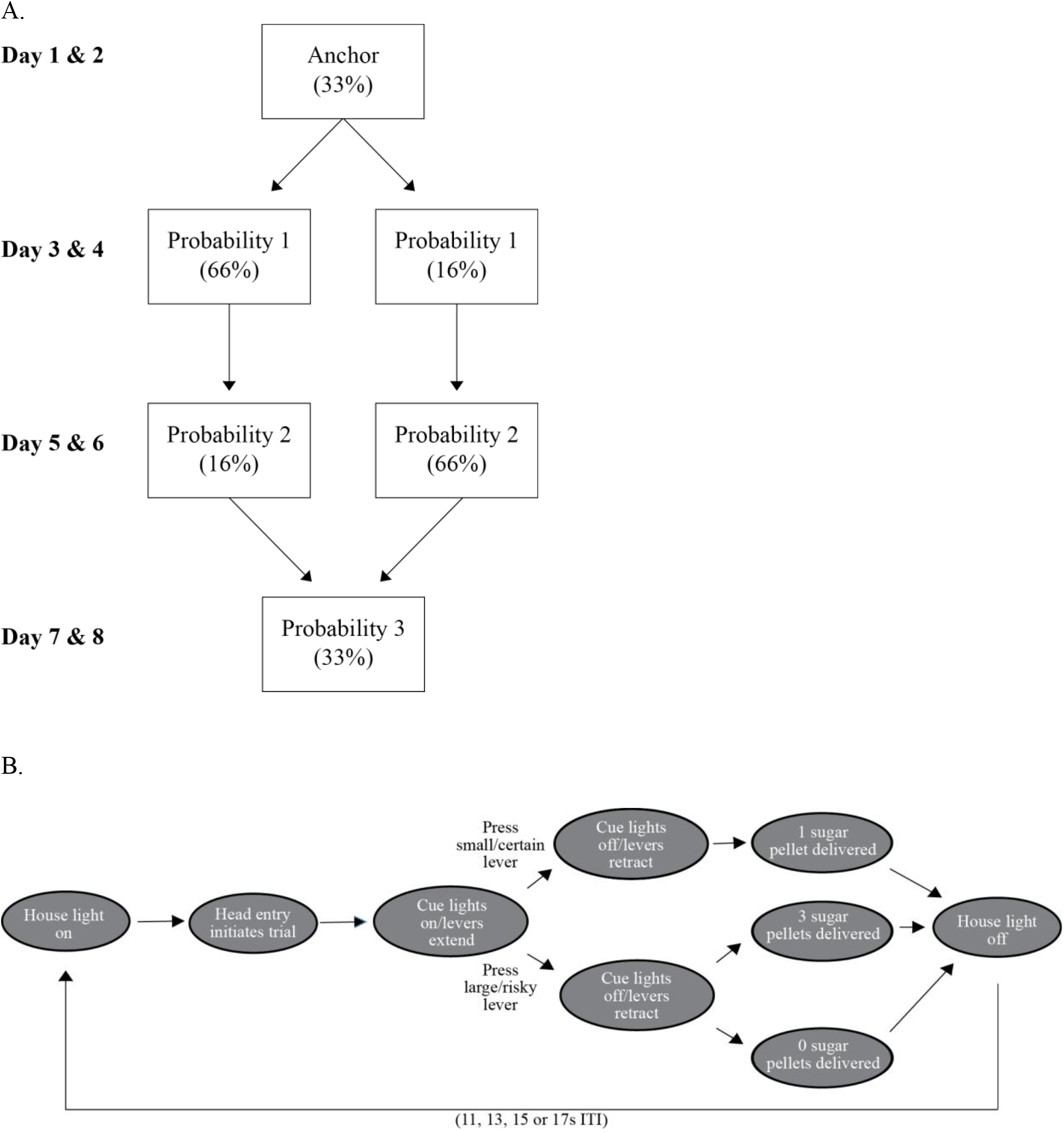
Risk task. (A) The order of probability sessions throughout the task was pseudo-randomized across animals. (B) The sequence of events during individual trials of magnitude discrimination, the probabilistic risk task, and extinction.

##### Extinction

Extinction sessions (large, risky lever payoff was 0%) were conducted following completion of the task.

##### Trial Structure

The opportunity for each trial began with the illumination of the house light. Trials were then initiated by the animal making a head entry into the central dispenser, which immediately generated the illumination of the cue light(s) for 2 seconds; this indicated which lever(s) would extend at the end of the cue period. After the lever(s) extended and the animal pressed one lever, the lever(s) retracted, the cue lights were extinguished, and a reward (or omission) was delivered 2 seconds after the lever press. Ten seconds after the reward (or omission) was dispensed, the house light was extinguished for a variable ITI (11, 13, 15, or 17s). The house light then illuminated again; animals were not able to initiate a new trial until the house light was illuminated (Figure 2B). Risk preference for each session was calculated as the percentage of large, risky lever presses during the free-choice block.

#### Lavage procedures

The estrous cycle of female rats was monitored daily during on days that rats performed the risk task. Samples were collected following completion of the behavior task by gently flushing the vaginal area using a disposable pipet filled with sterile water. Residual water containing cells was then examined on a glass slide under a microscope. Identification of estrous phase was based on the estimation of relative concentrations of leukocytes, nucleated epithelial cells, and cornified epithelial cells (Pompili, Tomaz, Arnone, Tavares, & Gasbarri, 2010).

#### Behavioral data analysis

A hierarchical linear regression was used to examine weight as a factor of gelatin condition, sex, and PD.

Consumption between animals was normalized by dividing the amount of gelatin consumed on any day by the animal’s weight that day (g/kg). Average alcohol and gelatin consumed was calculated as the average of all 2-hour standard access periods. A two-way ANOVA was used to examine the effect of sex and alcohol condition on average gelatin intake. A linear regression was used to test if gelatin consumed across standard access days could be explained by PD.

The relationship between BEL and amount of alcohol consumed on PD51 was examined with regression. Five samples were removed from the analysis because of contamination that was apparent upon visual inspection and affirmed with a BEL level that was less than 10 mg/dl. The cause of the contamination is unclear, it is possible that the heparin and blood did not mix properly before the plasma was extracted.

Behavior during sessions in which the outcome was certain (magnitude discrimination and extinction) were analyzed with a hierarchical linear regression using the predictors of probability of risky lever payoff, average alcohol consumed, and sex. Data used for this analysis included behavior from the last day of magnitude discrimination and extinction.

A two-way RM-ANOVA was used to analyze the effect of sex and probability of risky lever delivery on percent of risky lever presses in control animals. A hierarchical linear regression was used to further examine if there was an underlying correlation between sugar consumed in adolescence and risk preference in adulthood. A hierarchical linear regression was used to evaluate preference for the risky lever in animals that consumed alcohol in adolescence as predicted by probability of risky lever payoff, sex, and average alcohol consumed.

Further categorization of behavior employed quantification of win-stay decisions, lose-shift decisions, trials completed, and total pellets delivered. Separate linear regressions were used to examine trials completed and total pellets delivered over the session as a factor of sex, probability of risky lever payoff, and alcohol. A similar model was used to evaluate win-stay and lose-shift decisions; however, session half was incorporated as an independent variable. Win-stay decisions were defined as risky lever presses that immediately followed a risky lever press that had resulted in the delivery of 3 sugar pellets. Lose-shift decisions were identified as certain lever presses that immediately proceeded a risky lever press that resulted in the omission of sugar pellets.

A one-way RM-ANOVA was used to examine the effect of estrous cycle on risk preference.

### Electrophysiological Methods

#### Surgical procedures

Once animals completed behavior shaping (∼PD 77), they were anesthetized with ketamine-xylazine (0.1, 0.05 ml/kg, IP) and stainless-steel Teflon insulated electrode microwire arrays (MicroProbes, Gaithersburg, MD) were implanted bilaterally. The electrode arrays were organized into two rows of four microwires (50 μm diameter; tip separation .25mm) and were guided stereotaxically into mPFC (AP +3.3, ML +1.2 relative to bregma, and −3.5 mm relative to the brain’s surface at a 10° angle) and OFC (AP +3.2, ML −3.0 relative to bregma, and −4.0 mm relative to the brain’s surface). Surgical steel screws and dental cement were used to secure the implanted electrodes; ground wires from each array were secured around one of the distal surgical screws. Animals were given at least one week of recovery time after surgery before continuing onto magnitude discrimination.

#### Electrophysiological recordings

Electrophysiological recordings began when the rats were roughly PD 100 (Figure 1). Electrode arrays were connected to a recording cable that was attached to a motorized commutator (Plexon, Dallas, TX) allowing rats to move freely in the operant chamber. Electrical signals detected at the electrode tip were amplified and transduced via the OmniPlex system (Plexon, Dallas, TX). Events such as lever presentation, lever press, sugar pellet delivery, and nose port entry were time stamped onto the neural activity data using transistor-transistor logic (TTL). Individual waveform statistics were performed during the recording session to identify waveforms belonging to individual neurons (PlexControl). All recorded waveforms were evaluated again and refined after recordings had ended (Offline Sorter). The data were exported to Matlab and R for additional analysis.

#### Histology

When animals completed behavioral testing and extinction, the final electrode position was marked by passing electrical current down each electrode. The rats were euthanized, perfused, and the brains were removed and allowed to post-fix in a Potassium Ferricyanide, formalin mixture for 24 hours before being moved to a 20% sucrose solution. Brains were sliced at 40μm to assess electrode placement. Neurons were only included in the analyses if the wire they were recorded on had a verified placement in one of the regions of interest.

#### Electrophysiological data analysis

To determine whether each neuron showed a change in activity at the time of task events, its average time-course of the response was aligned to the time of lever press on each trial and the average firing rates during baseline (8-5 s before trial initiation), the time of the lever press (2s before lever press to 1s after lever press), and the reward period (2-8 s after lever press) were calculated. Neurons were excluded if they exhibited a baseline firing rate less than .3 sp/s or greater than 20 sp/s. Student’s t-tests were performed on each neuron to identify which exhibited a significant change in firing rate at the time of lever press or reward compared to baseline. Neurons were grouped based on event and direction of response and could be sorted into multiple groups. Firing rates were further sorted based on the lever that was pressed and the number of sugar pellets that resulted from the lever press.

To compare across neurons, each neuron’s mean firing rate and standard deviation of the firing rate over the entire session were calculated. Average firing rates were z-transformed using the neuron’s baseline mean and standard deviation firing rate. Hierarchical linear regressions were performed on each response group to test the relationship between z-transformed firing rate, sex, alcohol consumed in adolescence, and event (lever pressed or number of sugar pellets delivered). Analyses were repeated using more conservative degrees of freedom in a mixed model regression analysis, but the results were almost identical. Tables containing the more conservative results can be found in the Supplemental Results. Analyses were done separately in mPFC and OFC based on a priori hypotheses. At least one model for each neuron population tested the interaction between sex and alcohol; if the moderation was significant the sexes were analyzed separately to prevent the complicated interpretation of a three-way interaction. Neurons with z-scores above 10 were verified as outliers and excluded.

Independent one-way ANOVAs examined the effect of estrous cycle stage on baseline firing rate. Neurons were grouped based on their anatomical location and if on average they had a significant increase in firing rate at the time of lever press or reward, or a significant decrease at the time of lever press or reward.

R (version 3.4.3, R Core Team, 2017) and the R-packages *plyr* (Wickham, 2011)*, ggplot2* (Wickham, 2009)*, afex* (Singmann et al., 2018)*, lmerTest* (Kuznetsova et al., 2017)*, emmeans* (Lenth, 2018), and *car* (Fox & Weisberg, 2011) were used for analyses.

## RESULTS

### Weight

Access to alcohol did not affect the typical weight gain expected across adolescence in male or female animals (Figure 3). PD showed a predicted, positive effect on weight (Table 1, Model 1: b = 0.01, 95% CI [0.01, 0.01], t (479) = 50.39, p < 0.001) and explained 84% of the variance, (R^2^ = 0.84, F (1, 479) = 2539, p < 0.001). Sex was incorporated into Model 2 to test the hypothesis that it moderates the effect of PD on weight. There was an effect of sex, indicating females weighed less than males (b = −0.08, 95% CI [−0.10, −0.07], t (477) = −12.80, p < 0.001), and an interaction between PD and sex denoting that males had accelerated weight gain compared to females, (b = 0.002, 95% CI [0.002, 0.003], t (477) = 17.19, p < 0.001). Overall this improved the model fit (R^2^ = 0.95, F (3, 477) = 2868, p < 0.001). Finally, gelatin condition was added to the Model 3 to ensure that the alcohol condition had no effect on weight; this was confirmed by the results (b = −0.0002, 95% CI [−0.002, 0.002], t (476) = −0.21, p = 0.84). The interaction between sex and PD was quantified by examining the simple slopes of sex at −1 and +1 SD of PD. When rats were younger, male rats weighed more (b = −0.07, 95% CI [−0.08, −0.05], t (477) = −11.92, p < 0.001), but as they aged the effect of sex became stronger, and the weight discrepancy became larger (b = −0.1, 95% CI [−0.12, −0.09], t (477) = −13.43, p < 0.001).

**Figure 3.**
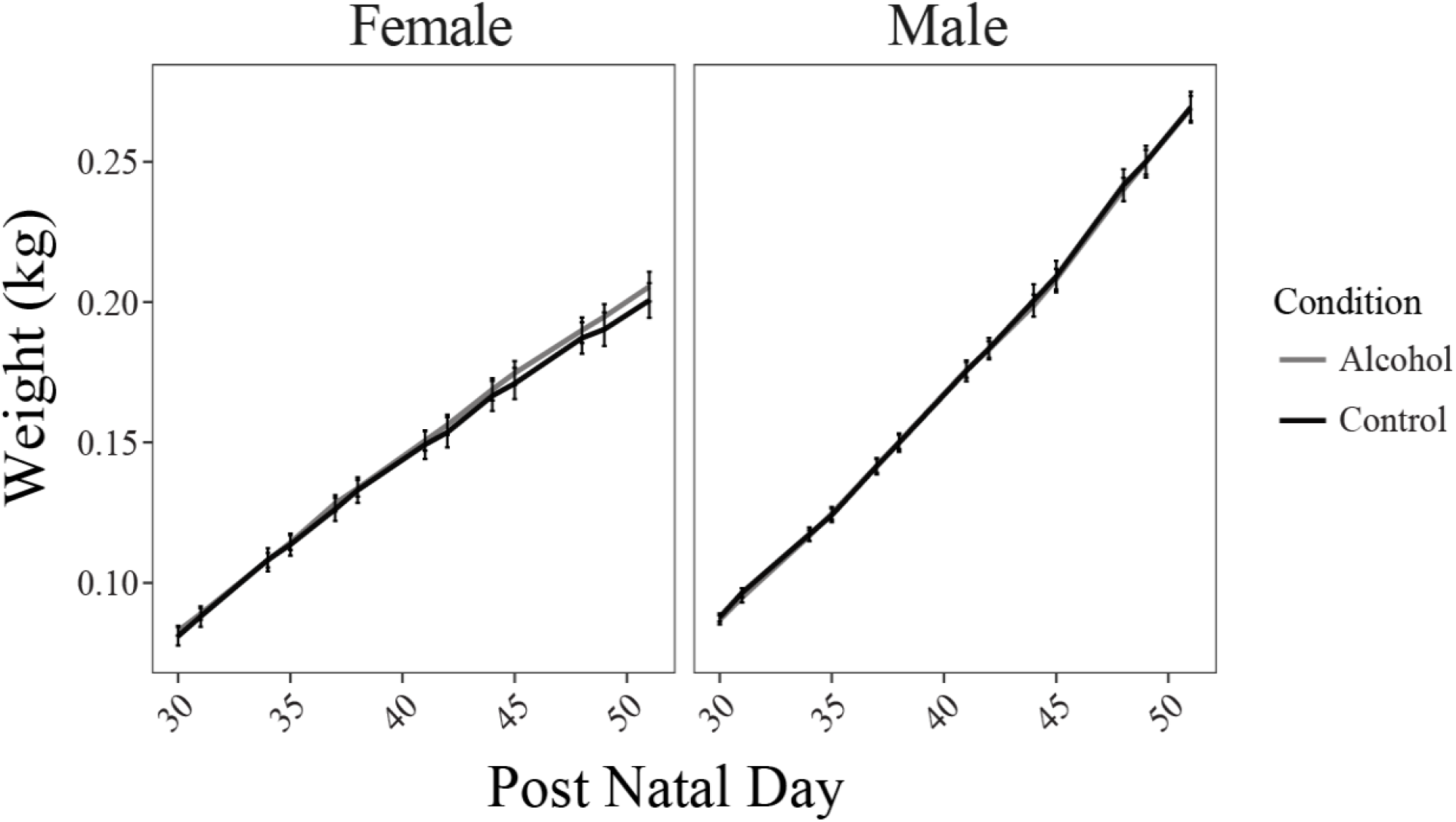
Weight of alcohol and control animals across all AIE days. Group is denoted by line color and error bars indicate SEM of each group by day.

**Table 1.** Summary of hierarchical regression analysis for variables predicting weight.

|  | Model 1 | Model 2 | Model 3 |
| --- | --- | --- | --- |
| Intercept | -0.133*** (0.0058) | -0.086*** (0.0050) | -0.085*** (0.0050) |
| PD | 0.007*** (0.0001) | 0.006*** (0.0001) | 0.006*** (0.0001) |
| Sex |  | -0.087*** (0.0068) | -0.087*** (0.007) |
| PD x Sex |  | 0.003*** (0.0002) | 0.003 *** (0.0002) |
| Condition |  |  | -0.0002 (0.001) |
| R<sup>2</sup> | 0.841 | 0.948 | 0.948 |
| F | 2539 | 2868 | 2147 |
| df<sub>1</sub> | 1 | 3 | 4 |
| df<sub>2</sub> | 479 | 477 | 476 |
| p | < 0.001 | < 0.001 | < 0.001 |

Note: \* p < 0.05, \*\* p < 0.01; \*\*\* p < 0.001

### Variable alcohol consumption

Gelatin consumption depended on whether or not it contained alcohol. There was a significant effect of gelatin condition on average amount of gelatin consumed (Figure 4A: F (1,33) = 110.19, p < 0.001) and a trend toward a main effect of sex (F (1,33) = 3.76, p =0.06), but there was no interaction between sex and gelatin condition on average gelatin consumed (Figure 4B: F (1, 33) = 0.15, p = 0.71). Least-squared means post-hoc comparisons revealed the control group (M = 43.44, SE = 2.07) on average, consumed significantly more than the alcohol group (M = 15.20, SE = 1.71). Males on average consumed 1.83 g of alcohol per kg body weight each day (M = 1.83, SD = 0.96) and females consumed 1.21 g of alcohol per kg body weight (M = 1.21, SD = 0.74).

**Figure 4.**
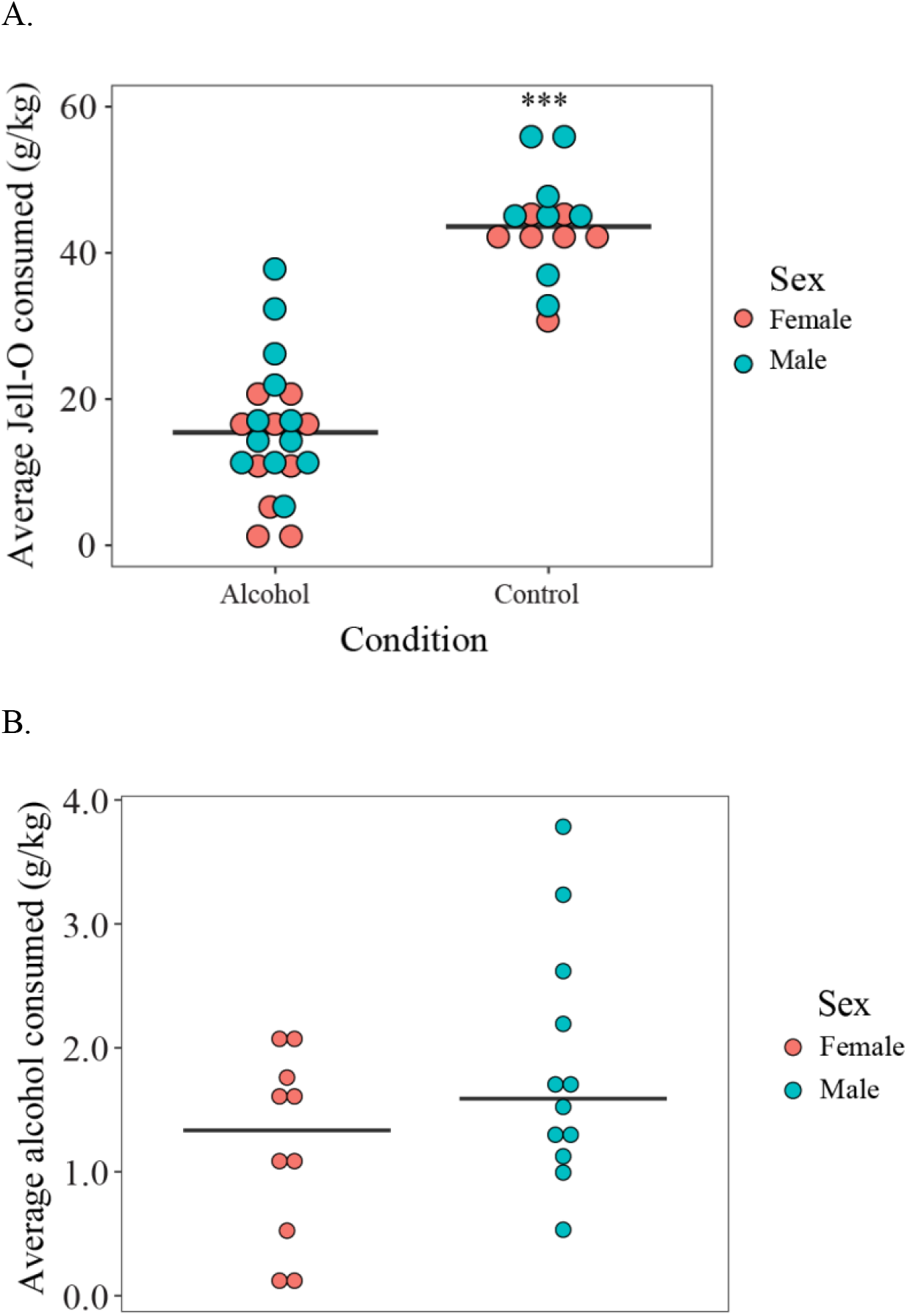
Average gelatin and alcohol consumption. (A) The average amount of gelatin consumed by individual rats across all 2-hour access days. Controls consumed significantly more gelatin than alcohol animals. (B) The average amount of alcohol consumed by individual rats in the alcohol group. Horizontal lines indicate the mean of each group (A) or sex (B). *** p < 0.001

Consumption of alcohol across the entire AIE period is shown separately for control and alcohol animals in Figure 5A,B. At the beginning of the AIE exposure, when access was provided for 24, 6, or 3 hours, animals consumed more gelatin than in the remaining days (Figure 5C,D). During the standard, 2-hour access days, PD did not have a significant effect on gelatin consumed (Figure 5E,F: b = 0.26, 95% CI [−0.15, 0.68], t (331) = 1.27, p = 0.20) and explained less than 1% of the variance (R^2^ = 0.005, F (1, 331) = 1.61, p = 0.20).

**Figure 5.**
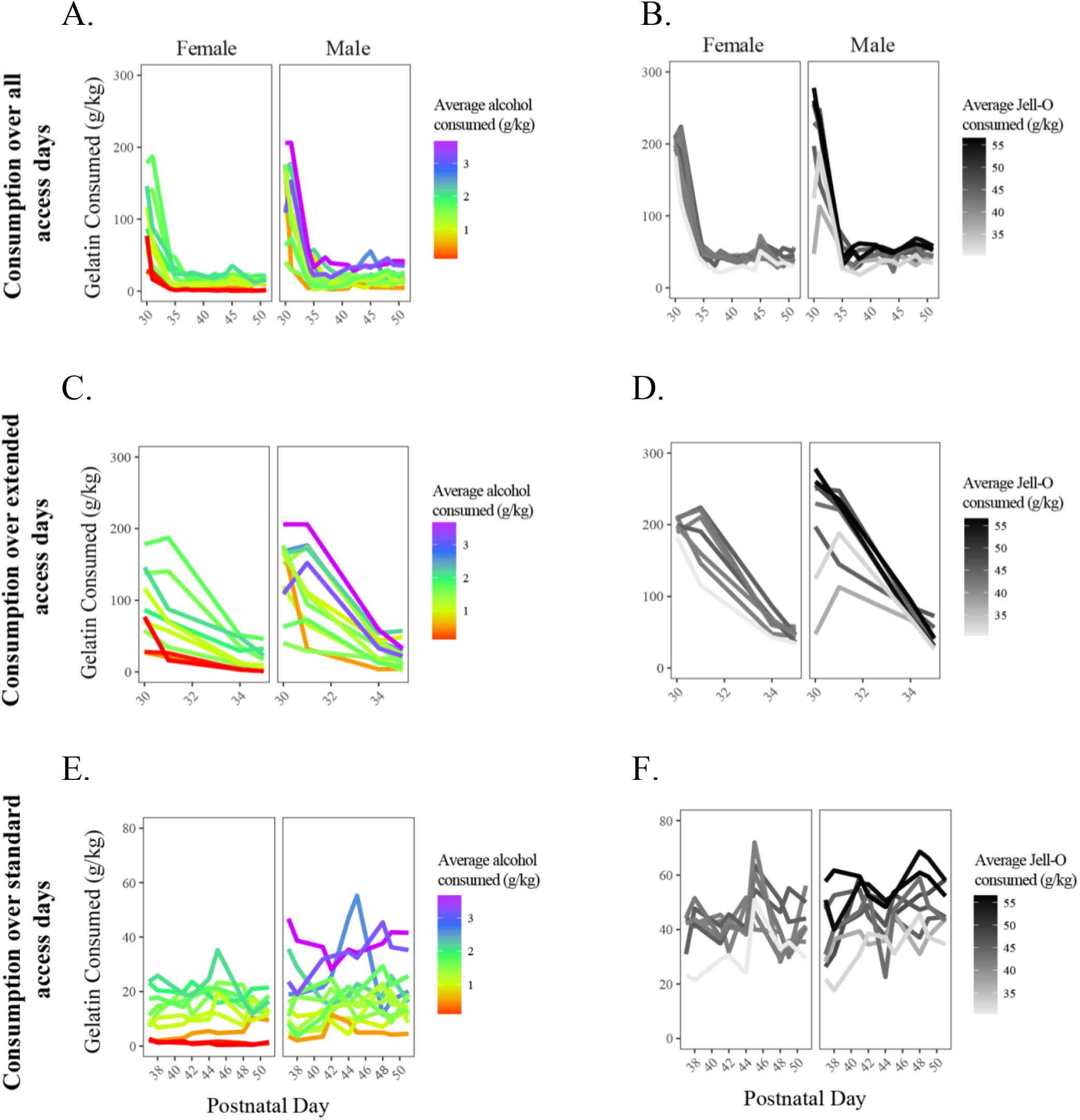
Patterns of individual gelatin consumption across AIE days. Lines indicate the behavior of individual rats and are colored based on the average alcohol consumed if they were in the alcohol group (A, C, E) or the average gelatin consumed if they were in the control group (B, D, F). Panel rows are based on patterns of gelatin consumption over all of adolescence (A, B), solely extended access days (C, D), or exclusively 2-hour standard access days (E, F).

### Blood ethanol levels

The best fit linear regression had a slope of 18.93, such that for every 1g/kg alcohol, blood ethanol increased by 18.93mg/dl.

### Risk task performance

When there was no uncertainty about reward delivery or omission - during magnitude discrimination and extinction phases of testing (100% and 0% chance of large, risky reward delivery, respectively)- rats reliably chose the lever with the larger reward option, regardless of prior alcohol consumption (Figure 6). We examined the percentage of large, risky lever choices during magnitude discrimination and extinction as a factor of probability of risky lever payoff, sex, and average alcohol consumed in adolescence (Table 2, Model 1). This model resulted in a significant, positive effect of probability (b = 0.76, 95% CI [0.72, 0.80], t (70) = 33.90, p < 0.001) such that magnitude discrimination elicited more risky lever presses than extinction. There was no effect of sex (b = −0.03, 95% CI [−0.05, − 0.01], t (70) = −1.19, p = .24) or average alcohol (b = 0.02, 95% CI [0.01, 0.03], t (70) = 1.92, p = 0.06) on risk preference. Model 1 accounted for 94% of the variability in the dependent variable, (R^2^ = 0.94, F (3, 70) = 384.5, p < 0.001). We also examined if there was an interaction between any of the predictors in Model 1. This strengthened the effect of probability (Table 2, Model 2: b = 0.91, 95% CI [0.81, 1.07], t (66) = 9.73, p <0.001) and there was a significant moderation of sex on probability (b = −0.13, 95% CI [− 0.18, −0.07], t (66) = −2.14, p = 0.04). Although Model 2 did improve the R^2^ fit by 0.01 it was not a significantly better fit than Model 1 (F (1,66) = 2.48, p = 0.052).

**Figure 6.**
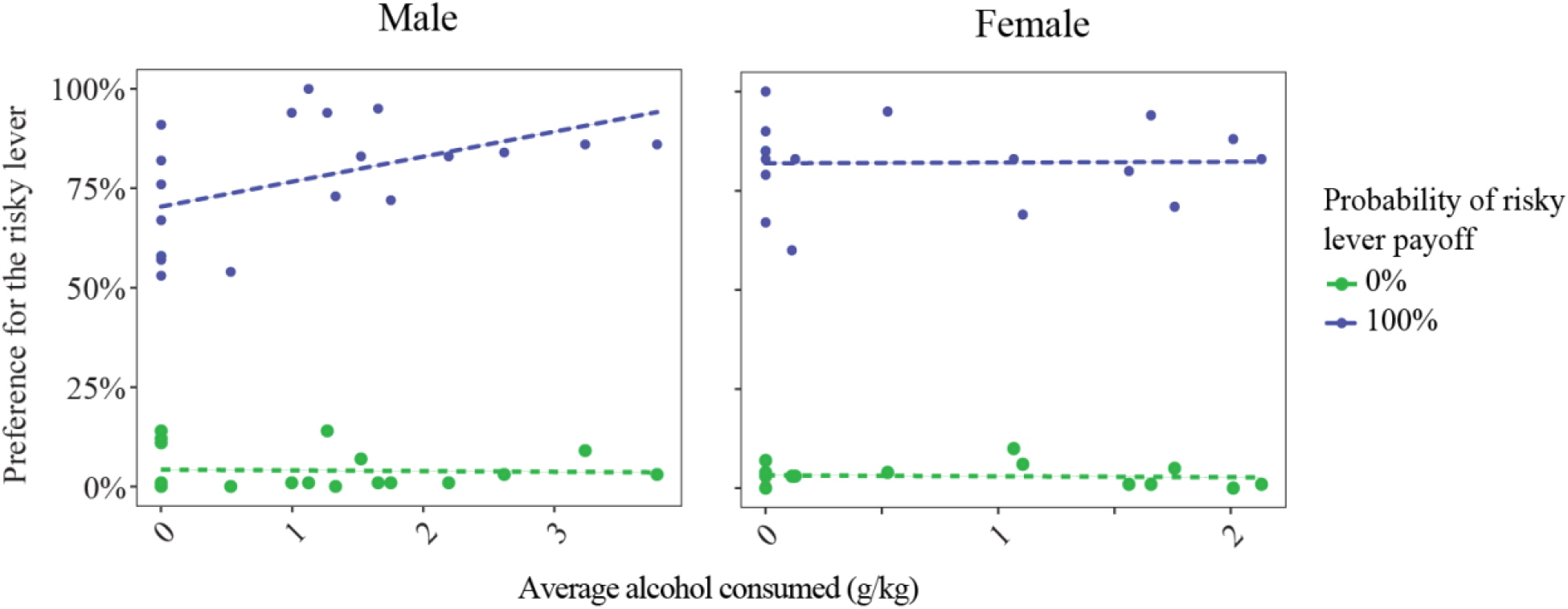
Preference for the risky lever during magnitude discrimination and extinction. Dots denote individual rat’s percentage of large/risky lever presses based on the average amount of alcohol they consumed in adolescence. Lines represent the simple slopes of each probability session across average alcohol consumption. Dots and lines are colored based on the probability of the risky lever payoff.

**Table 2.**
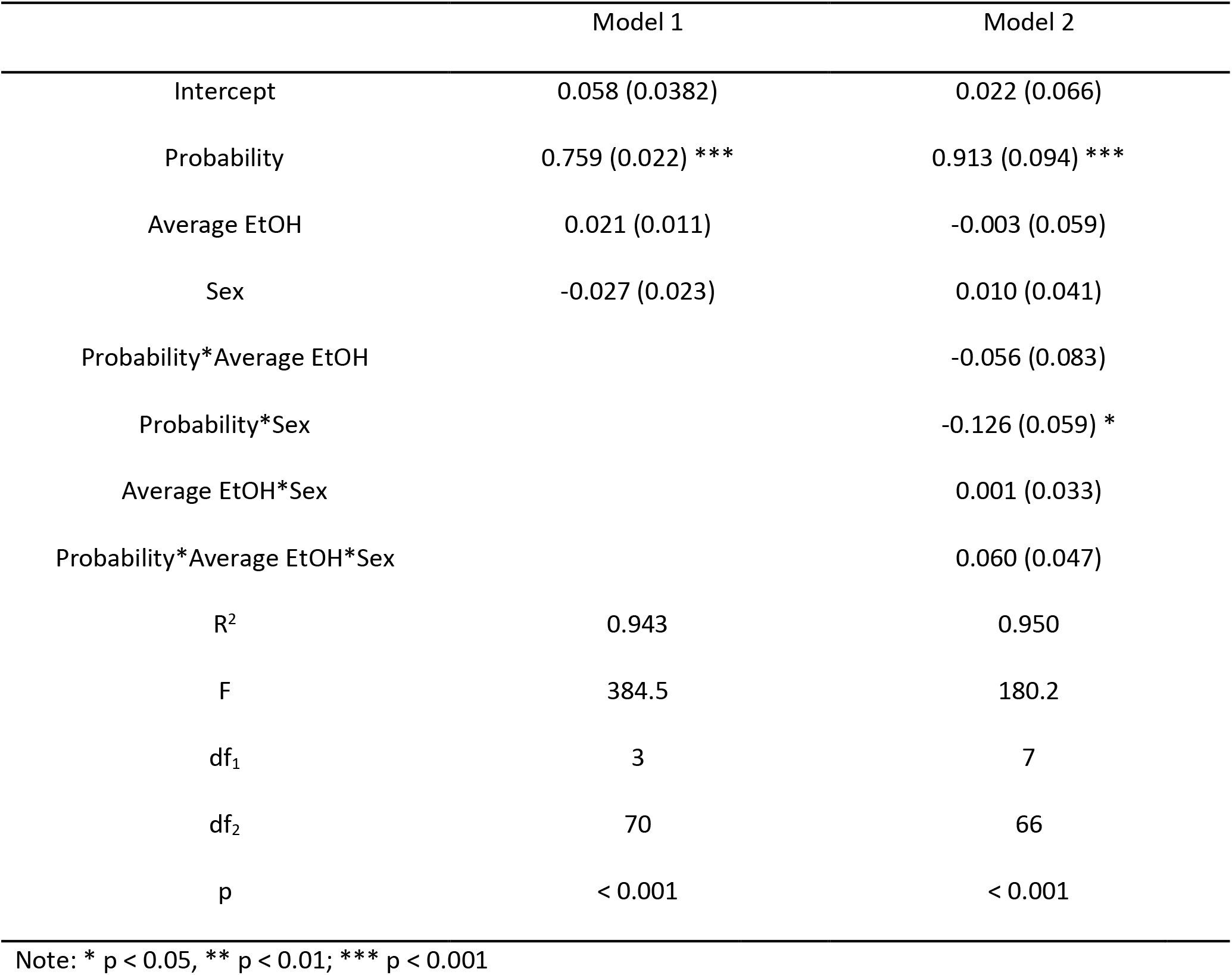
Summary of linear regression analysis for variables predicting large/risky lever presses during magnitude discrimination and extinction.

|  | Model 1 | Model 2 |
| --- | --- | --- |
| Intercept | 0.058 (0.0382) | 0.022 (0.066) |
| Probability | 0.759 (0.022) *** | 0.913 (0.094) *** |
| Average EtOH | 0.021 (0.011) | -0.003 (0.059) |
| Sex | -0.027 (0.023) | 0.010 (0.041) |
| Probability*Average EtOH |  | -0.056 (0.083) |
| Probability*Sex |  | -0.126 (0.059) * |
| Average EtOH*Sex |  | 0.001 (0.033) |
| Probability*Average EtOH*Sex |  | 0.060 (0.047) |
| R<sup>2</sup> | 0.943 | 0.950 |
| F | 384.5 | 180.2 |
| df<sub>1</sub> | 3 | 7 |
| df<sub>2</sub> | 70 | 66 |
| p | < 0.001 | < 0.001 |

Note: \* p < 0.05, \*\* p < 0.01, \*\*\* p < 0.001

In control animals, preference for the risky lever depended on the probability of large reward, but did not differ between sexes or depend on the amount of gelatin consumed. During performance of the risk task (16%, 33%, and 66% chance of large, risky reward), control animals showed a significant main effect of probability of risky lever payout on selection of the risky lever indicating that they pressed the lever more as its probability of payout increased (Figure 7A: F (2,28) = 15.33, p < 0.001). However, there was no effect of sex on lever preference (F (1,13) = 0.18, p = 0.68). Post-hoc tests using least-squared means revealed no difference between 16% (M = 0.39, SE = 0.15) and 33% (M = 0.51, SE = 0.16) or between 33% and 66% (M = 0.74, SE = 0.16), but there was a significant difference between 16% and 66%. A linear regression, Model 1 in Table 3, confirmed probability of risky lever payout was a strong predictor of preference for the risky lever (b = 0.71, 95% CI [0.55, 0.87], t (42) = 4.35, p <0.001), but sex was not (b = 0.04, 95% CI [−0.01, 0.09], t (42) = 0.58, p = 0.57). Overall Model 1 accounted for 31% of the variability in the dependent variable (R^2^ = 0.31, F (2, 42) = 9.61, p < 0.001). Average gelatin consumed in adolescence was incorporated into Model 2 (Figure 7B, Table 3) but this did not have an effect on risk preference in adulthood (b = 0.01, 95% CI [0.00, 0.02], t (41) = 1.25, p = 0.22), nor did it increase the explained variance (R^2^ = 0.34, F (3, 41) = 7.007, p < 0.001), (F (1) = 1.56, p = 0.22).

**Figure 7.**
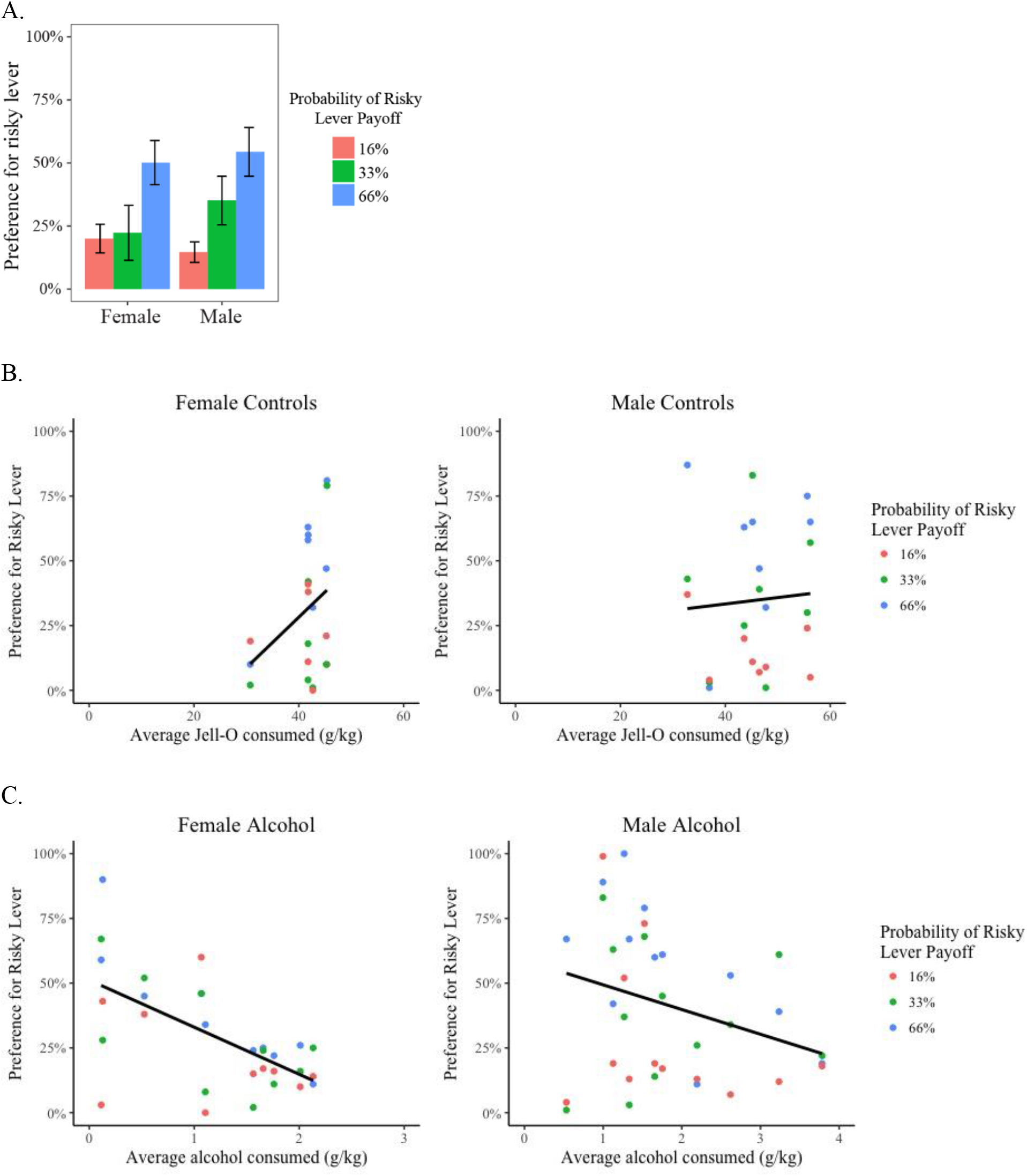
Effect of adolescent alcohol use on probabilistic risk task performance. (A) Average risky lever preference of control animals by risky reward probability. Error bars indicate SEM. (B) The percentage of risky lever presses by control animals across the range of gelatin consumed in adolescence. (C) The percentage of risky lever presses by alcohol animals across the range of alcohol consumed in adolescence. Dots indicate the behavior of individuals and dot color is representative of the session probability and the black lines denote the linear regression fit of gelatin (B) or alcohol (C) as a predictor of preference for the risky lever for each sex.

**Table 3.** Summary of hierarchical regression analysis for variables predicting large/risky lever presses in control rats.

|  | Model 1 | Model 2 |
| --- | --- | --- |
| Intercept | -0.001 (0.125) | -0.245 (0.231) |
| Probability | 0.705*** (0.162) | 0.705*** (0.0.161) |
| Sex | 0.039 (0.068) | 0.011 (0.071) |
| Average gelatin |  | 0.007 (0.005) |
| R<sup>2</sup> | 0.314 | 0.225 |
| F | 9.605 | 7.007 |
| df<sub>1</sub> | 2 | 3 |
| df<sub>2</sub> | 42 | 41 |
| p | <0.001 | <0.001 |

Note: \* p < 0.05, \*\* p < 0.01; \*\*\* p < 0.001

In animals that consumed alcohol in adolescence, preference for the risky lever depended on the probability of large reward, sex, and the amount of gelatin consumed. The effects of probability of risky lever payoff and sex on preference for the risky lever in animals that had consumed alcohol in adolescence was examined in Model 1 (see Table 4). This model showed a significant effect of the probability of the risky lever (b = 0.46, 95% CI [0.32, 0.60], t (63) = 3.18, p = 0.002) such that preference for the lever increased with probability of lever payoff. In addition, males selected the risky lever more often than females (b = 0.12, 95% CI [0.06, 0.18], t (63) = 2.01, p = 0.005). Together these variables explained a significant portion of the variance in preference for the risky lever (R^2^ = 0.18, F (2, 63) = 7.08, p = 0.002). Model 2 incorporated average alcohol consumed to test the effect of adolescent alcohol use on percentage of risky lever choices. Surprisingly, increased alcohol use was associated with a reduction in preference for the risky lever. (Figure 7C, b = −0.12, 95% CI [−0.15, −0.09], t (62) = −3.75, p <0.001). The effect of probability of payout (b = 0.46, 95% CI [0.43, 0.49], t (62) = 3.49, p < 0.001) and sex remained (b = 0.20, 95% CI [0.14, 0.26], t (62) = 3.40, p = 0.001). Model 2 increased explained variance to 33% (R^2^ = 0.33, F (3, 62) = 10.4, p < 0.001) and ANOVA results show Model 2 was a significantly better fit than Model 1 (F (1) = 14.09, p < 0.001). Thus, preference for the risky lever increased with increasing reward probability, was higher in males, but decreased with higher levels of adolescent alcohol consumption.

**Table 4.** Summary of hierarchical regression analysis for variables predicting large/risky lever presses in alcohol rats.

|  | Model 1 | Model 2 |
| --- | --- | --- |
| Intercept | -0.006 (0.113) | 0.066 (0.104) |
| Probability | 0.461** (0.145) | 0.461*** (0.001) |
| Sex | 0.122* (0.060) | 0.200** (0.059) |
| Average EtOH |  | -0.124*** (0.033) |
| R<sup>2</sup> | 0.184 | 0.335 |
| F | 7.081 | 10.40 |
| df<sub>1</sub> | 2 | 3 |
| df<sub>2</sub> | 63 | 62 |
| p | 0.002 | <0.001 |

Note: \* $p < 0.05$ , \*\* $p < 0.01$ , \*\*\* $p < 0.001$

To examine the effect of prior trial outcomes on choices, we analyzed patterns of win-stay/lose-shift behavior across individual sessions. The relationship between win-stay choices between the first and second half of the test session is shown separately for males and females for control animals and those who had access to alcohol in Figure 8A. Win-stay decisions was evaluated by the factors sex, alcohol, probability of risky lever payoff, and session half which collectively explained 35% of variability (R^2^ = 0.35, F (4, 177) = 23.78, p < 0.001). Males engaged in significantly more win-stay decisions than females (b = −2.34, 95% CI [−2.97, −1.71], t (177) = −3.71, p = 0.003). Additionally, win-stay decisions decreased with increased adolescent alcohol use (b = −0.70, 95% CI [−1.00, −0.04], t (177) = −2.33, p = 0.02) and increased as the probability of the risky lever increased (b = 0.12, 95% CI [0.11, 0.13], t (177) = −4.34, p < 0.001). Finally, more of win-stay decisions occurred in the second half of the session compared to the first (b = 1.38, 95% CI [0.79, 1.97], t (177) = 2.33, p = 0.02).

**Figure 8.**
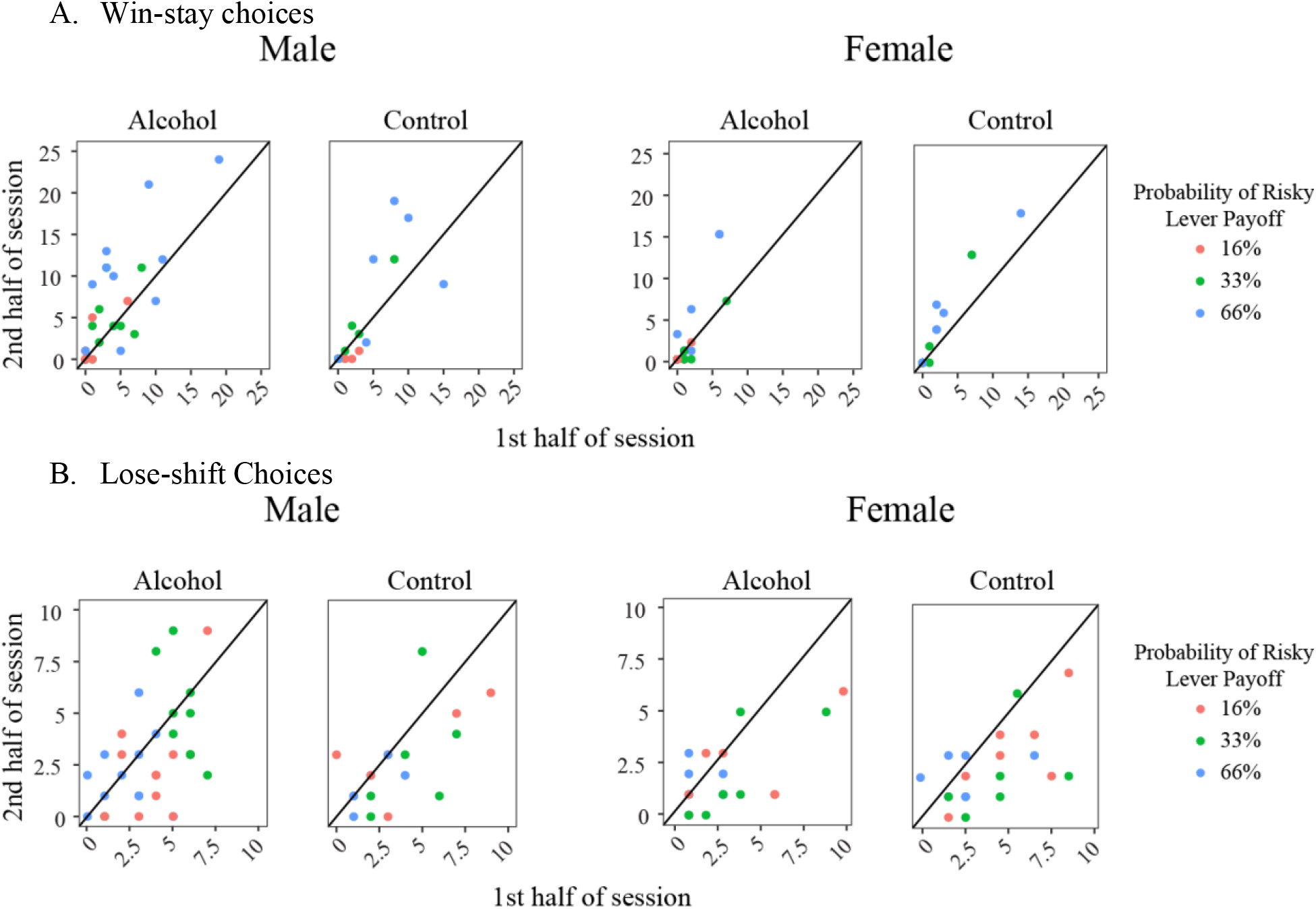
Win-stay and lose-shift decisions indicate ongoing adjustments in decision-making behavior. (A) A comparison of the number of win-stay decisions in the first half of the session vs the 2^nd^ half in males and females across groups. (B) A comparison of the number of lose-shift decisions in the first half of the session vs the 2^nd^ half as a factor of sex and group. Dots represent the behavior of individuals and are colored based on the probability of the risky lever payoff of the session. Black lines represent the line of unity, such that points that fall below indicate higher frequency in the first half of the sessions and above indicate higher frequency in the second half.

The relationship between lose-shift choices between the first and second half of the test session is shown separately for males and females for control animals and those who had access to alcohol in Figure 8B. As the probability of risky lever payoff increased, the number of lose-shift choices decreased (b = −0.03, 95% CI [−0.04, −0.03], t (177) = −4.38, p < 0.001) and more lose-shift decisions were made in the first half of the session than the second (b = −1.00, 95% CI [−1.32, −0.68], t (177) = −3.15, p = 0.002). However, sex (b = −0.45, 95% CI [−0.78, 0.11], t 177) = −1.34, p = .18) and adolescent alcohol use (b = 0.07, 95% CI [−0.09, 0.23], t (177) = 0.43, p = 0.66) did not significantly contribute to the variability in lose-shit decisions (R^2^ = 0.15, F (4, 177) = 7.95, p < 0.001). Overall, the behavioral patterns suggest that when animals experienced reward omission early in the sessions, they were more likely to switch to the certain lever and maintain that choice pattern.

To assess general task engagement, we considered the total number of trials completed in sessions and total number of sugar pellets earned. Overall, males completed more trials per session than females, and fewer trials were completed as the probability of the risky lever increased (Figure 9A). The number of trials completed was significantly predicted by sex (b = −9.93, 95% CI [−13.37, −6.49], t (88) = − 2.89, p = 0.005) and probability of risky lever payoff (b = −0.16, 95% CI [−0.24, −0.08], t (88) = −2.08, p = 0.04). Adolescent alcohol use did not significantly contribute (b = 0.16, 95% CI [−0.66, 0.97], t (88) = 0.19, p = 0.85) to the explained variance in the model (R^2^ = 0.14, F (3, 88) = 4.71, p = 0.004). Similarly, males received more sugar pellets than females (Figure 9B: b = −12, 95% CI [−17.06, −8.45], t (88) = −2.96, p = 0.004). Probability of risky lever payoff was also a significant, positive factor of total sugar pellets received (b = 0.92, 95% CI [0.82, 1.02], t (88) = 9.46, p < 0.001). Again, alcohol consumed in adolescence was not related to total sugar pellets delivered (b = −0.50, 95% CI [−1.53, 0.53], t (88) = −0.48, p = 0.63). Overall sex, probability of risky lever payout and alcohol explained 53% of the variance (R^2^ = 0.53, F (3, 88) = 32.9, p < 0.001).

**Figure 9.**
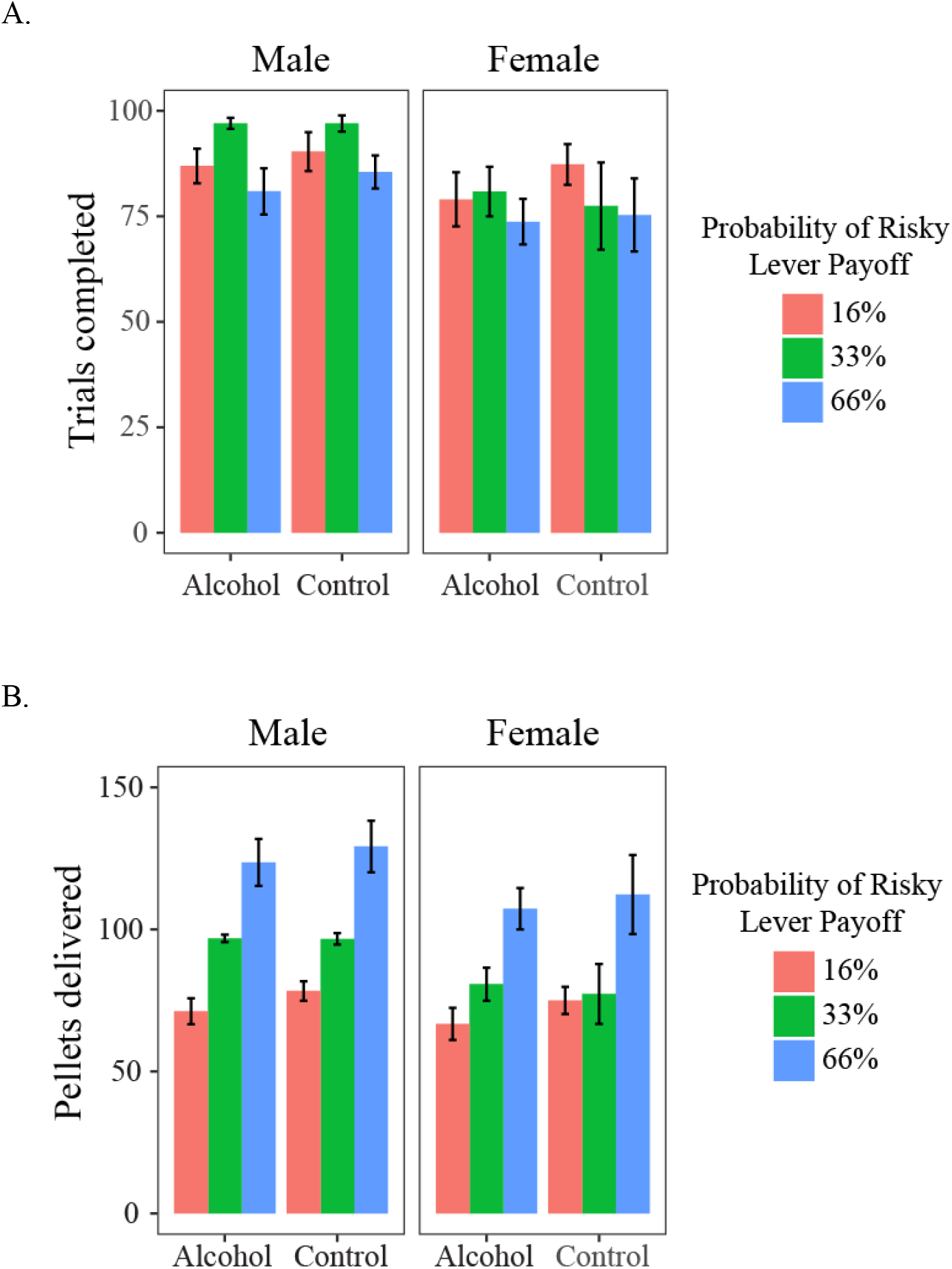
The number of trials completed and total pellets delivered are indicative of task engagement. (A) Number of trials completed as a factor of sex and group. (B) Total pellets received over a session as a factor of sex and group. Bar color indicates the probability of risky lever reward delivery, error bars denote SEM.

### Estrous cycle

There was no effect of estrous cycle on preference for the risky lever (F (3,53) = 0.55, p = 0.65; Figure 10). Female rats exhibited a normal estrous cycle that lasted between 3-5 days.

**Figure 10.**
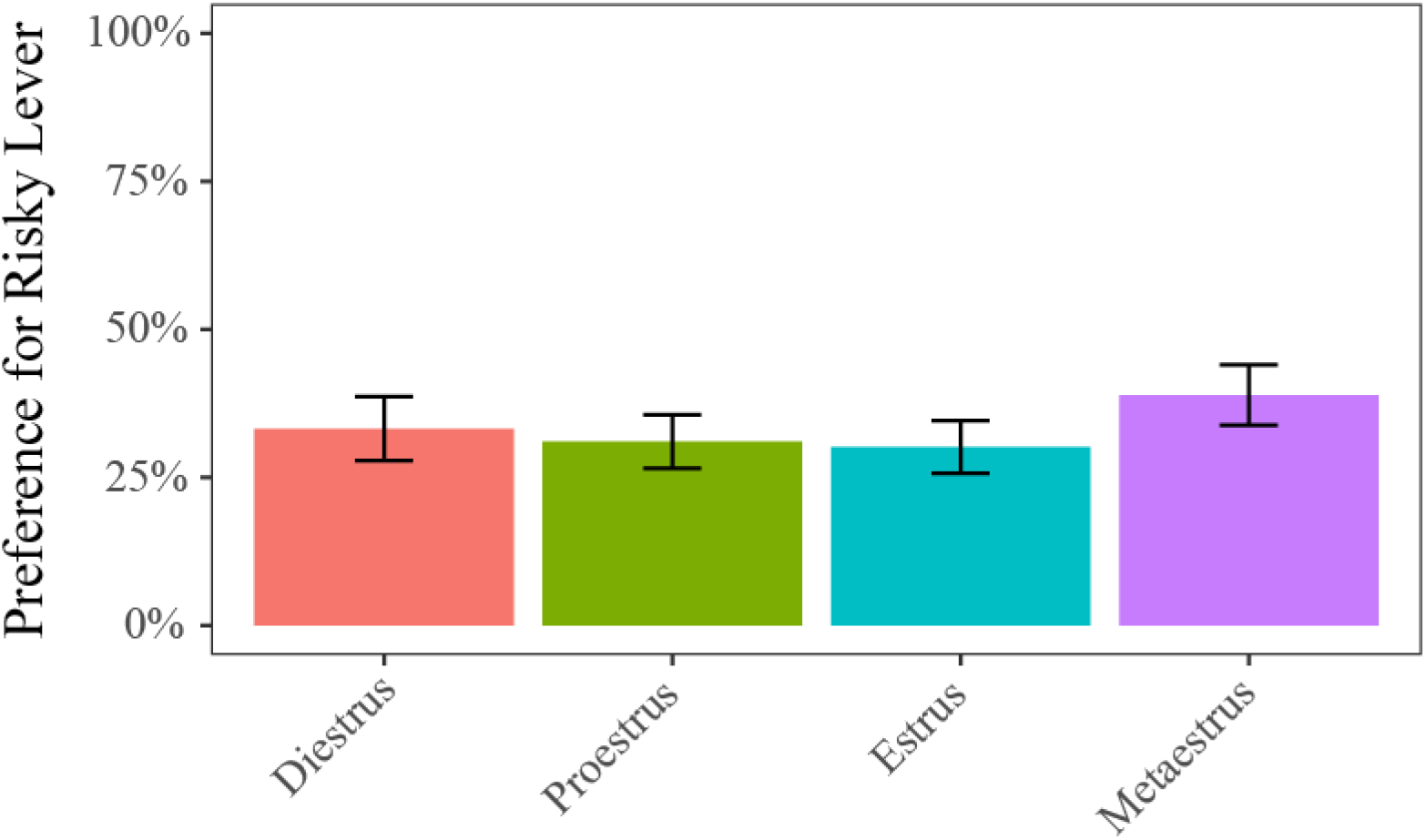
Effect of estrous cycle on probabilistic risk task performance. Estrous cycle stage was not a predictor of female rats’ performance on the probabilistic risk task. Error bars indicate SEM.

### Time course of responses and classification

In total, we recorded from 1,186 neurons; 60 were excluded due to electrode misplacement (15) or poor isolation with too few actions potentials identified (45). Electrode placements can be seen in Figure 11. Of the remaining 1126 neurons, 493 of these neurons were recorded in mPFC and 633 were recorded from OFC. 578 of the neurons came from males and 548 came from females. Control animals contributed 482 neurons to the total, while 644 neurons were in alcohol animals.

**Figure 11.**
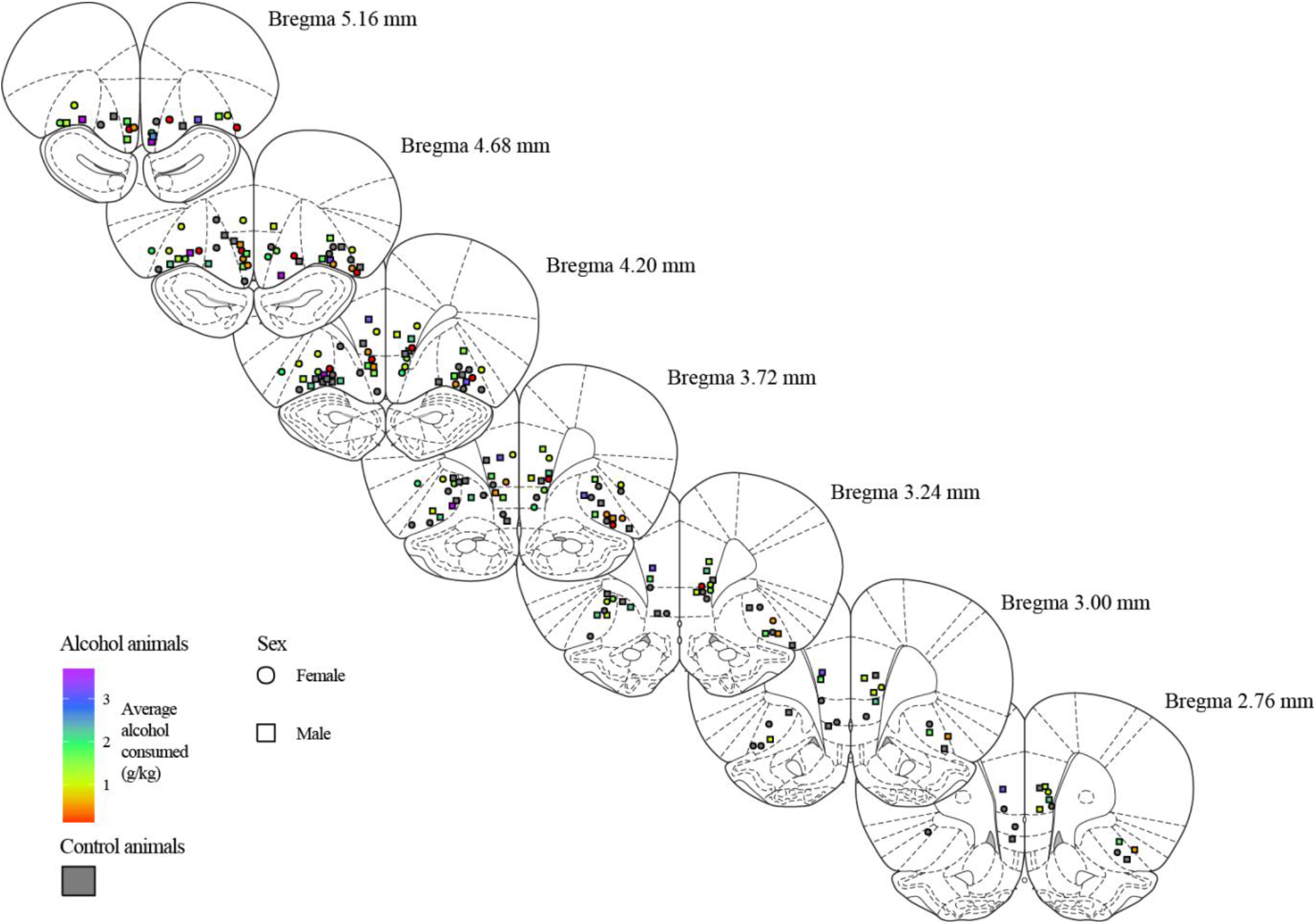
Histologically confirmed electrode placement in OFC and mPFC. The shape of label indicates what sex the wire was in and color indicates the average amount of alcohol consumed in adolescence.

We first looked at the time-course of all neurons in response to trial events. The illumination of the house light did not have an effect on the average baseline firing rate of neurons (2s before house light, M = 2.45, SD = 0.07 sp/s; 2s after house light, M = 2.54, SD = 0.07 sp/s; t (1125) = −0.87, p = 0.38) (Figure 12A). Aligning activity to the onset of the trial (i.e. nosepoke into central port), we observed a small response to the illumination of the cue light(s), and a larger response 2s later following lever presentation (Figure 12B). At the time of lever presentation, it is not known which lever the rat will select and which outcome will be delivered. To analyze the effects of choice and outcome on neural responses, we focused on the timing of activity aligned to lever press (Figure 12C). The epochs of interest were baseline activity (5-8 s before the press), press-related activity (2 s before to 1 s after the press), and reward delivery (or omission; 2 to 8 s after the press).

**Figure 12.**
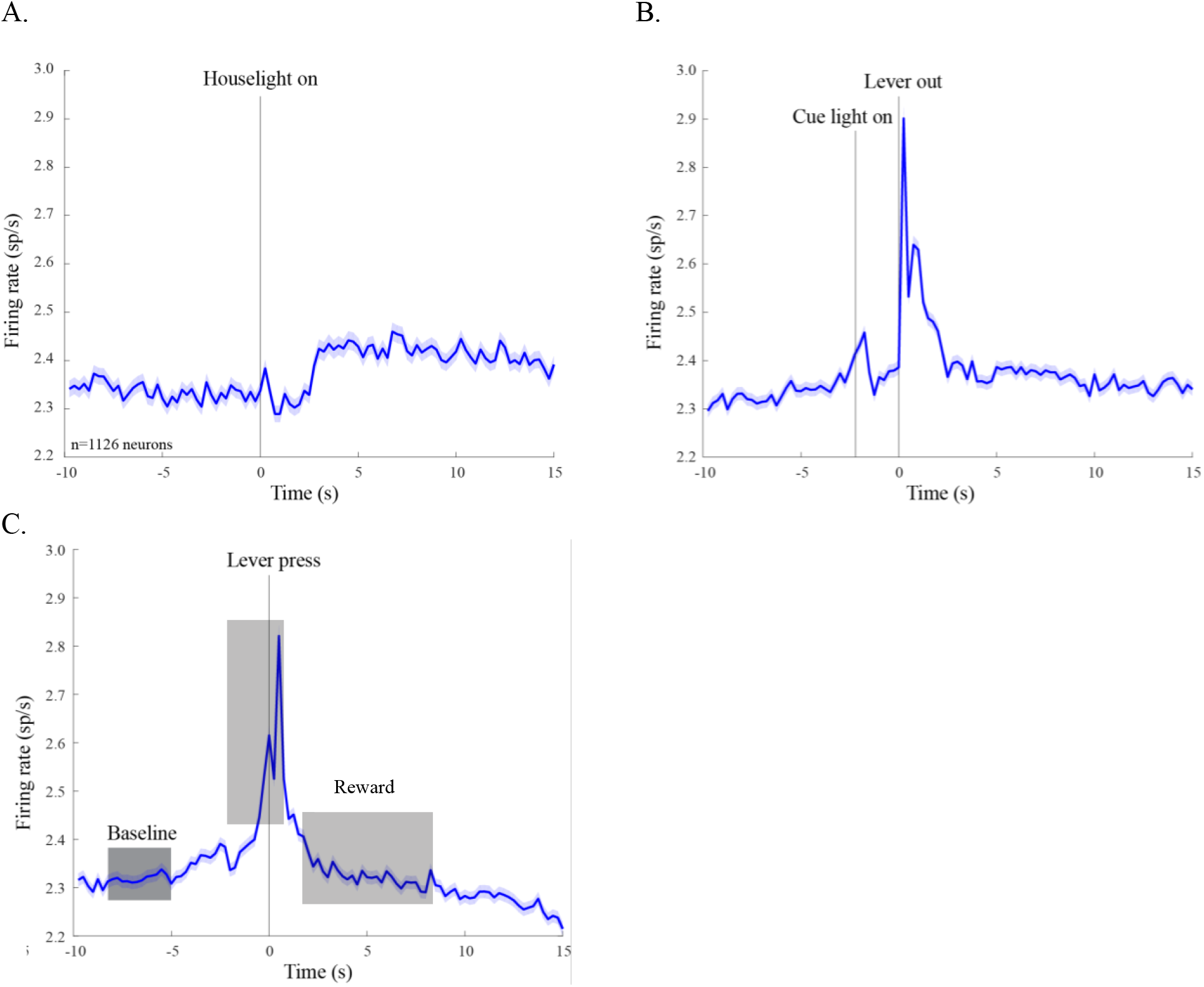
Average activity of all neurons aligned to external stimuli. (A) Neural activity aligned to the illumination of the house-light reveals no effect of the house-light on baseline activity. (B) Activity aligned to the extension of the levers shows a brief increase in activity at the time of trial initiation, denoted as the time the cue lights turned on. (C) Activity aligned to the time of lever press shows earlier environmental stimuli did not cause any fluctuations in baseline firing rate. Epochs used in subsequent analyses are shaded in gray.

Neurons were grouped into 4 categories based on their average patterns of activity: increased activity at lever press (PP), decreased activity at lever press (NP), increased activity at reward delivery (PR) or decrease in activity at reward delivery (NR). mPFC contained 108 PP neurons, 126 NP neurons, 115 PR neurons, and 163 NR neurons. OFC contained 139 PP neurons, 104 NP neurons, 173 PR neurons, and 152 NR neurons (Table 5). mPFC and OFC contained different distributions of neurons as confirmed by a chi-squared analysis (χ^2^ = 15.20, p = 0.002). The time course of neural activity in mPFC for each group (PP, NP, PR, NR) aligned to the time of lever press is shown separately for controls and alcohol-consuming animals in Figure 13. The time course of activity for PP, NP and NR neurons in OFC is shown in Figure 14. Because there were sex differences in OFC PR neurons (discussed below), their responses are shown separately for males and females in Figure 15.

**Figure 13.**
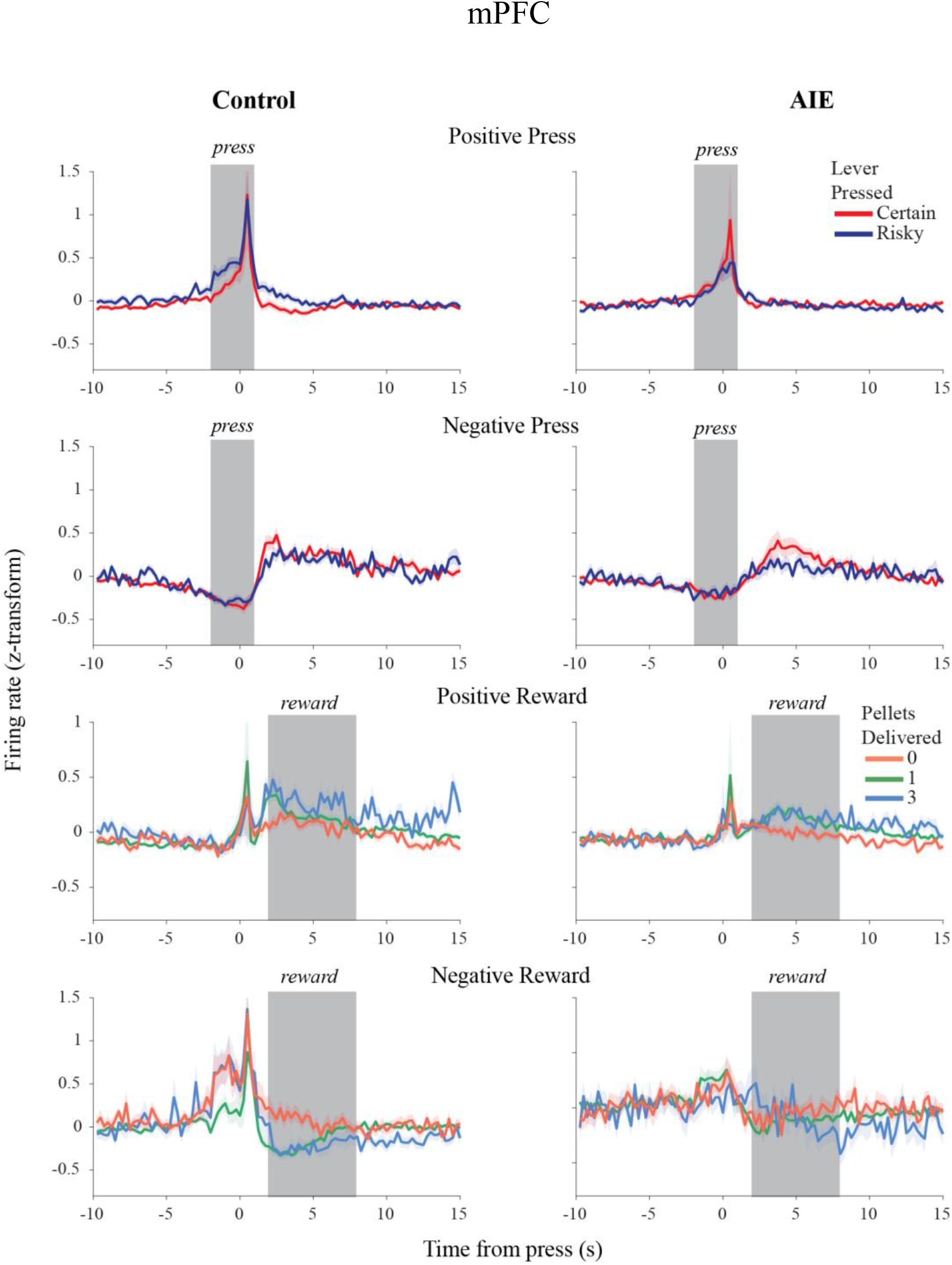
Time course of average neural activity in of four response types (PP, NP, PR, NR) in mPFC. Averages were calculated in 250ms bins aligned to time of lever press. Gray shaded regions indicate period of response that was compared to baseline. Neurons from control and AIE animals are presented separately. Color of line indicates the lever acted on or the outcome of the trial.

**Figure 14.**
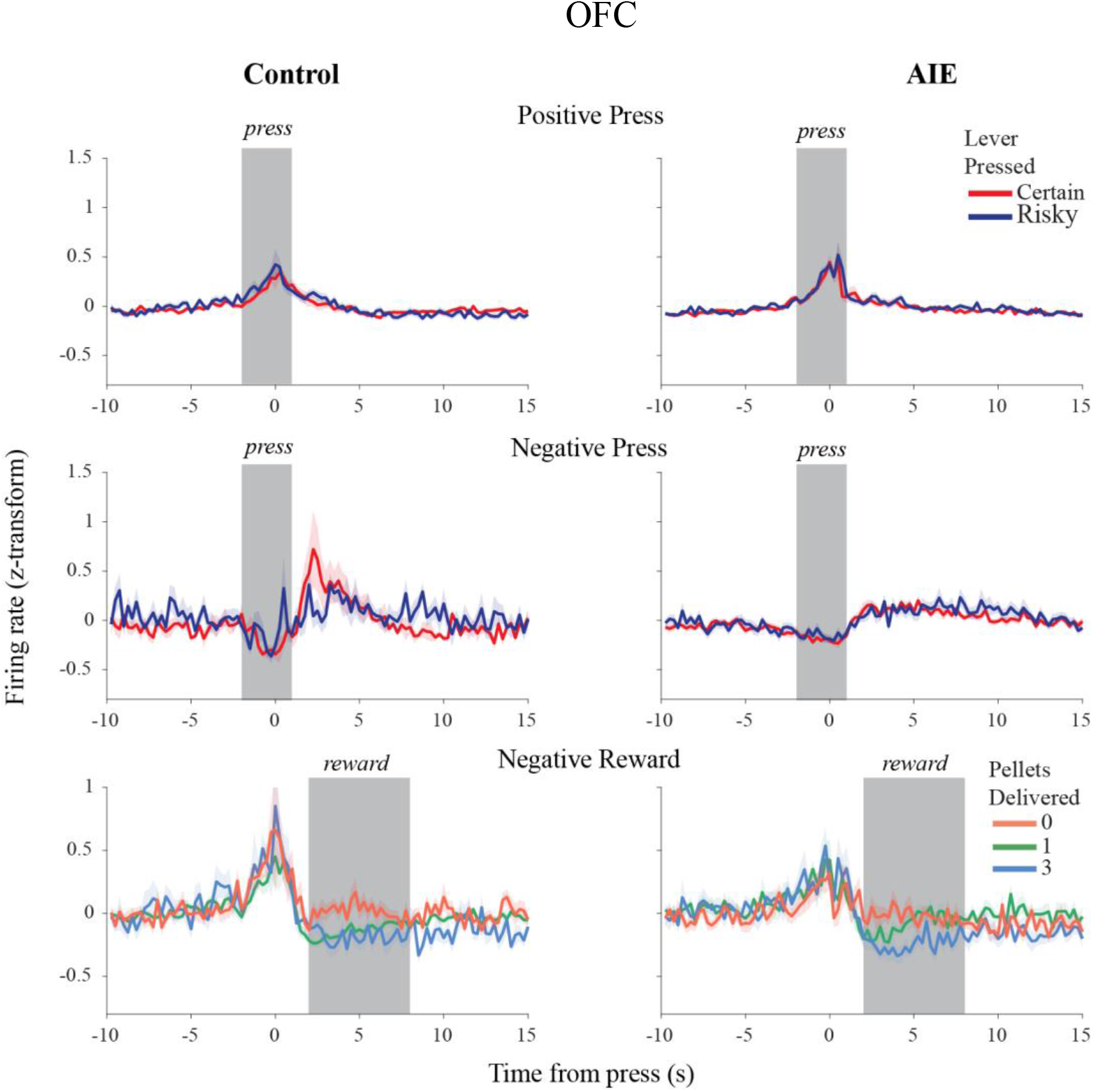
Time course of average activity in of PP, NP, NR neurons in OFC. Averages were calculated in 250ms bins aligned to time of lever press. Same conventions as Figure 13.

**Figure 15.**
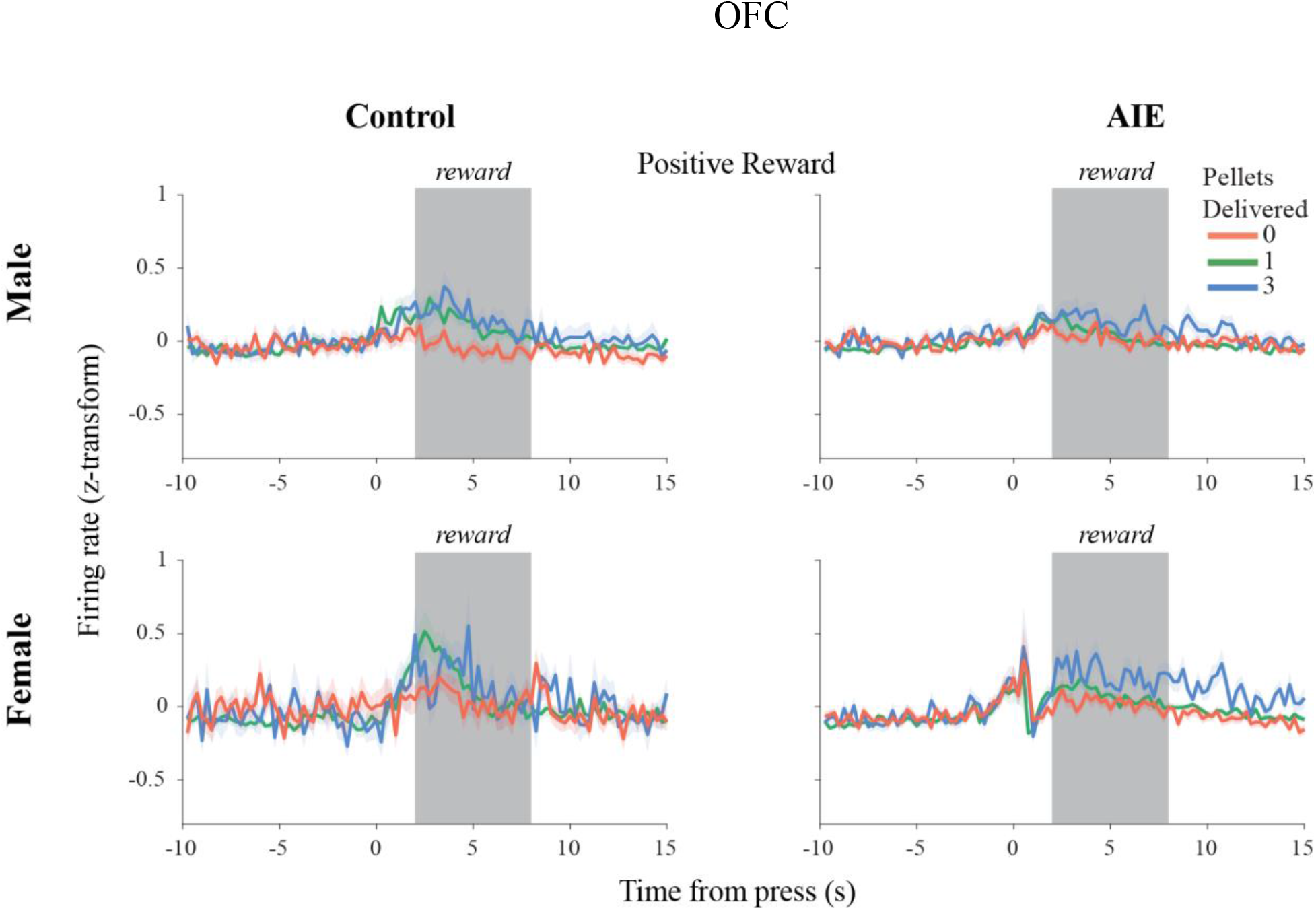
Time course of average neural activity of response type PR in OFC. Averages were calculated in 250ms bins aligned to time of lever press. Same conventions as Figure 13.

**Table 5.** Summary of neuron distributions in mPFC and OFC.

|  | mPFC | OFC |
| --- | --- | --- |
| Increase in activity during press<br>(PP) | 108 | 139 |
| Decrease in activity during press<br>(NP) | 126 | 104 |
| Increase in activity reward<br>delivery (PR) | 115 | 173 |
| Decrease in activity reward<br>delivery (NR) | 163 | 152 |

### Neural activity in mPFC

Subpopulations of mPFC neurons had activity that was modulated at the time of the lever press or during the period of reward delivery. The average responses of mPFC PP neurons at the time of lever press are shown as a function of alcohol consumption in Figure 16A. We first examined the activity in mPFC PP neurons as a function of sex, alcohol, and lever pressed (Table 6, Model PPm1). Females showed a higher change in firing rate from baseline than males (b = 0.39, 95% CI [0.22, 0.55], t (212) = 2.38, p = 0.02). There was a trend towards the risky lever eliciting a greater change in firing rate than the certain lever, but it was not significant (b = 0.28, 95% CI [0.13, 0.42], t (212) = 1.90, p = 0.06). There was no main effect of alcohol on change in firing rate during the time of lever press (b = −0.09, 95% CI [−0.19, 0.01], t (212) = −0.93, p = 0.35). Model PPm1 accounted for 6% of the variability in the firing rate at the time of lever press, (R^2^ = 0.06, F (3, 212) = 4.65, p = 0.004). Model PPm2 tested the interaction between sex and alcohol and confirmed that the effect of alcohol was similar between sexes (b = −0.18, 95% CI [−0.38, 0.02], t (211) = −0.91, p = 0.36), directing the continuation of the analysis without the interaction term. Model PPm3 tested if there was an interaction between lever pressed and alcohol, as it was hypothesized that neurons would show increased activity to the risky lever with increased adolescent alcohol use. There was a trend towards a significant moderation that contradicted our hypothesis (b = − 0.32, 95% CI [−0.49, −0.25], t (211) = −1.87, p = 0.06), such that increased alcohol use was correlated with a decrease in firing during risky lever presses and an increase in firing during certain lever presses. The addition of this interaction term also increased the beta coefficient and significance of the lever press term (b = 0.48 95% CI [0.30, 0.66], t (211) = 2.65, p = 0.009). Model PPm3 increased the explained variance by 2% (R^2^ = 0.08, F (4, 211) = 4.40, p = 0.002), and although it was not a significant improvement from Model PPm1 (F (1, 211) = 3.90, p = 0.06), it trended toward being a better fit, suggesting Model PPm3 is the best fit model for mPFC’s PP subpopulation’s neural activity.

**Figure 16.**
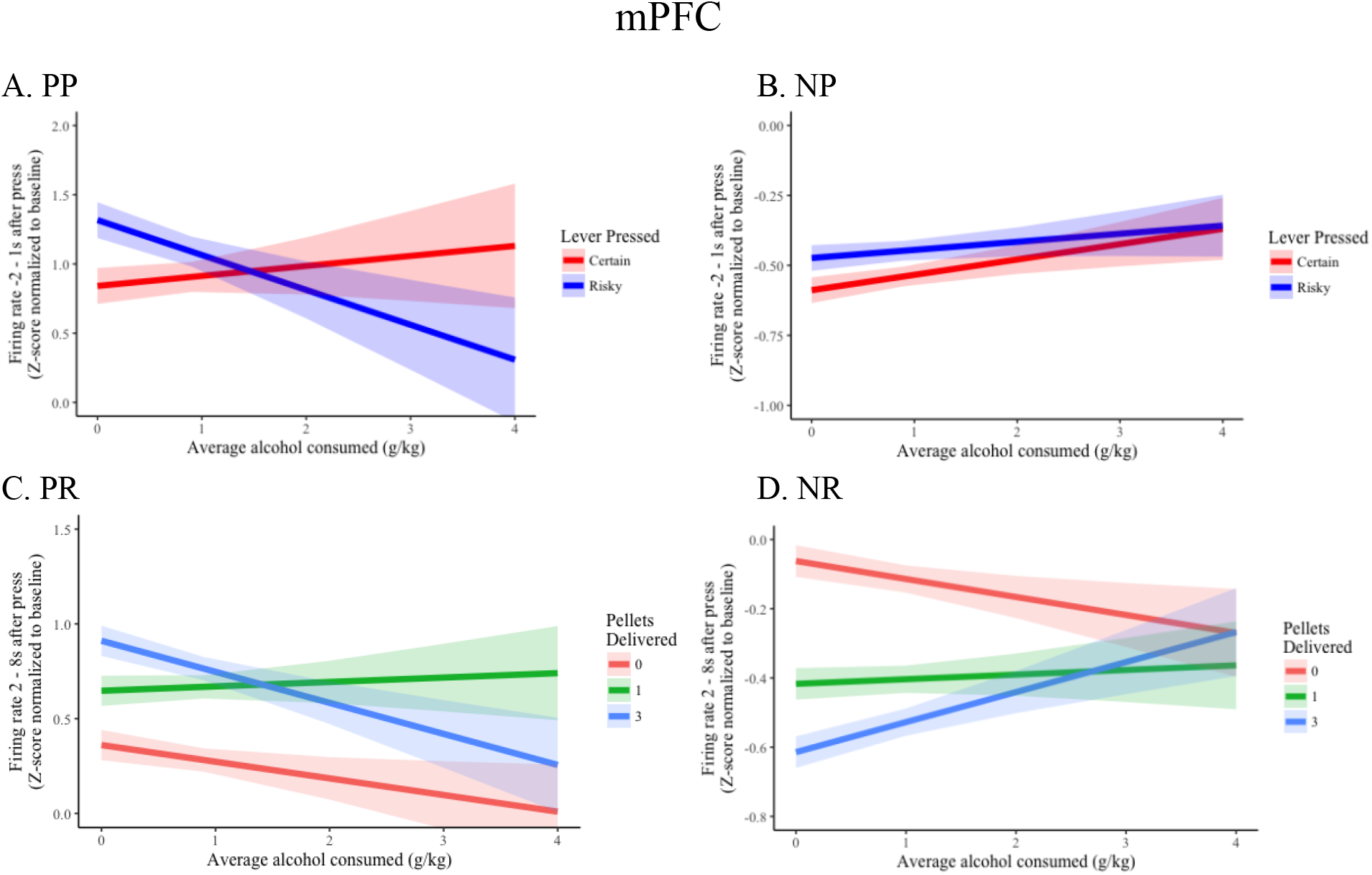
mPFC responses. Simple slopes of chosen lever, moderated by alcohol consumed in adolescence, predicting firing rate at the time of lever press in PP population (A) and NP population (B). Simple slopes of reward size, moderated by alcohol consumed in adolescence, predicting firing rate during the reward period in PR population (C) and NR population (D).

**Table 6.** Summary of hierarchical regression analysis for variables predicting change in firing rate during lever press in **mPFC PP** neurons.

|  | Model PPm1 | Model PPm2 | Model PPm3 |
| --- | --- | --- | --- |
| Intercept | 0.723*** (0.164) | 0.648*** (0.183) | 0.623*** (0.172) |
| Sex | 0.386* (0.161) | 0.49* (0.200) | 0.386* (0.160) |
| Average EtOH | -0.090 (0.097) | -0.016 (0.126) | 0.072 (0.130) |
| Lever Pressed | 0.276^ (0.145) | 0.276^ (0.145) | 0.476** (0.180) |
| Average EtOH*Sex |  | -0.179 (0.196) |  |
| Average EtOH*Lever Pressed |  |  | -0.325^ (0.174) |
| R<sup>2</sup> | 0.062 | 0.065 | 0.077 |
| F | 4.651 | 3.694 | 4.398 |
| df<sub>1</sub> | 3 | 4 | 4 |
| df<sub>2</sub> | 212 | 211 | 211 |
| p | 0.004 | 0.006 | 0.002 |

Note: ^ p < 0.1 \* p < 0.05, \*\* p < 0.01; \*\*\* p < 0.001

The average responses of mPFC NP neurons at the time of lever press are shown as a function of alcohol consumption in Figure 16B. We sought to model the neural activity in mPFC’s NP subpopulation of neurons, at the time of lever press, as a function of sex, alcohol, and lever pressed (Table 7, Model NPm1). This did not reveal a main effect of sex (b = 0.02, 95% CI [−0.04, 0.08], t (248) = 0.31, p = 0.76) or alcohol (b =0.04, 95% CI [0.01, 0.07, t (248) = 1.64, p = 0.10). There was a trend towards the risky lever generating a greater increase in firing compared to the certain lever (b = 0.09, 95% CI [0.04, 0.14], t (248) = 1.86, p = 0.06). Overall, Model NPm1only explained 3% of the variance and was not a significant predictor of change in firing rate at lever press (R^2^ = 0.03, F (3, 248) = 2.18, p = 0.09). Similar to Model PPm1, Model NPm2 confirmed that there was no interaction between sex and adolescent alcohol use (b =0.08, 95% CI [0.01, 0.15], t (247) = 1.17, p = 0.24). Finally, Model NPm3 tested if alcohol moderated the effect of lever. The beta coefficients for the other variables remained constant, but Model NPm3 did not yield a significant interaction (b = −0.03, 95% CI [−0.07, 0.01], t (248) = −0.59, p = 0.55) and the correlation coefficient did not increase (R^2^ = 0.03, F (4, 247) = 1.72, p = 0.15). Overall the decrease of NP neurons was stereotyped across sex, alcohol consumed, and lever chosen.

**Table 7.**
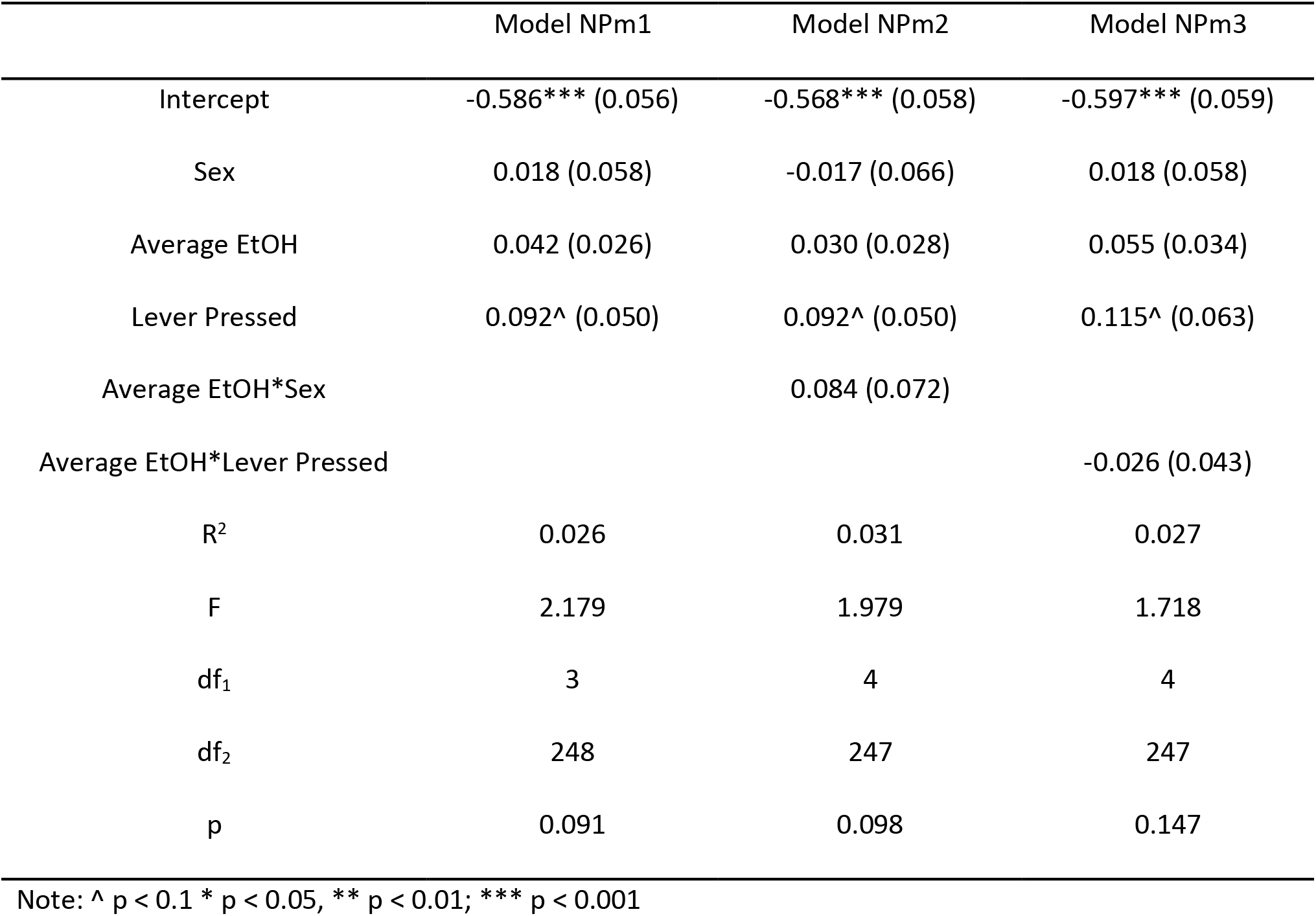
Summary of hierarchical regression analysis for variables predicting change in firing rate during lever press in **mPFC NP** neurons.

|  | Model NPm1 | Model NPm2 | Model NPm3 |
| --- | --- | --- | --- |
| Intercept | -0.586*** (0.056) | -0.568*** (0.058) | -0.597*** (0.059) |
| Sex | 0.018 (0.058) | -0.017 (0.066) | 0.018 (0.058) |
| Average EtOH | 0.042 (0.026) | 0.030 (0.028) | 0.055 (0.034) |
| Lever Pressed | 0.092^ (0.050) | 0.092^ (0.050) | 0.115^ (0.063) |
| Average EtOH*Sex |  | 0.084 (0.072) |  |
| Average EtOH*Lever Pressed |  |  | -0.026 (0.043) |
| R<sup>2</sup> | 0.026 | 0.031 | 0.027 |
| F | 2.179 | 1.979 | 1.718 |
| df<sub>1</sub> | 3 | 4 | 4 |
| df<sub>2</sub> | 248 | 247 | 247 |
| p | 0.091 | 0.098 | 0.147 |

Note: ^ p < 0.1 \* p < 0.05, \*\* p < 0.01; \*\*\* p < 0.001

For mPFC PR neurons, the increase in activity depended on the size of reward delivered, which was marginally moderated by adolescent alcohol consumption (Figure 16C). The activity of mPFC’s PR subpopulation during the reward period was first examined for effects of sex, adolescent alcohol use, and reward size (Table 8, Model PRm1). The number of sugar pellets delivered had a positive relationship with firing rate such that 1 sugar pellet increased normalized firing at the time of reward by 0.37 (b = 0.37, 95% CI [0.29, 0.45], t (340) = 4.29, p < 0.001) and 3 sugar pellets increased the firing rate by 0.50 (b = 0.50, 95% CI [0.41, 0.59], t (340) = 5.81, p < 0.001) compared to 0 pellets. However, the increases in firing rate following delivery 1 or 3 sugar pellets did not differ (b = 0.13, 95% CI [0.04, 0.22], t (340) = 1.53, p = 0.13). There was a trend toward a significant, negative relationship between alcohol and change in firing rate (b = −0.08, 95% CI [−0.13, −0.03], t (340) = −1.66, p = 0.10). In total, Model PRm1 explained 10% of the variance in firing rate (R^2^ = 0.10, F (4, 340) = 1.72, p < 0.001). The interaction between sex and alcohol was examined in Model PRm2, but it was not significant (b = 0.06, 95% CI [− 0.03, 0.15], t (339) = 0.626, p = 0.53) and the overall model was not significantly different than Model PRm1 (F (1, 339) =0.39, p = 0.53). Lastly, Model PRm3 examined if alcohol moderated the effect of reward size on firing rate during the reward period. There was no difference between alcohol’s effect on reward when comparing 0 and 1 pellets (b = 0.11, 95% CI [0.01, 0.21], t (338) = 1.11, p = 0.27) or 0 and 3 pellets (b = −0.76, 95% CI [−0.18, 0.02], t (338) = −7.61, p = 0.45), but the comparison between 1 pellet and 3 pellets verged on significance (b = −0.19, 95% CI [−0.29, −0.09], t (338) = −1.875, p = 0.06), such that the response to 3 pellets delivered compared to 1 weakened with higher alcohol consumption. Overall, Model PRm3 only explained 1% more variance in the dependent variable (R^2^ = 0.11, F (6, 338) = 7.14, p < 0.001) than Model PR1 and was not statistically different (F (2,338) = 1.78, p = 0.17).

**Table 8.** Summary of hierarchical regression analysis for variables predicting change in firing rate during the reward period in **mPFC PR** neurons.

|  | Model PRm1 | Model PRm2 | Model PRm3 |
| --- | --- | --- | --- |
| Intercept | 0.388*** (0.089) | 0.417*** (0.100) | 0.397*** (0.097) |
| Sex | -0.066 (0.078) | -0.107 (0.102) | -0.066 (0.078) |
| Average EtOH | -0.076 (0.046) | -0.101^ (0.061) | -0.088 (0.074) |
| Reward 0 v Reward 1 | 0.366*** (0.085) | 0.366*** (0.085) | 0.286* (0.111) |
| Reward 0 v Reward 3 | 0.496*** (0.085) | 0.496*** (0.085) | 0.550*** (0.111) |
| Reward 1 v Reward 3 | 0.130 (0.085) | 0.130 (0.085) | 0.264 * (0.111) |
| Average EtOH*Sex |  | 0.058 (0.092) |  |
| Average EtOH*Reward 0 v 1 |  |  | 0.111 (0.998) |
| Average EtOH*Reward 0 v 3 |  |  | -0.076 (0.998) |
| Average EtOH*Reward 1 v 3 |  |  | -0.187^ (0.100) |
| R<sup>2</sup> | 0.103 | 0.104 | 0.112 |
| F | 9.769 | 7.879 | 7.135 |
| df<sub>1</sub> | 4 | 5 | 6 |
| df<sub>2</sub> | 340 | 399 | 338 |
| p | < 0.001 | < 0.001 | < 0.001 |

Note: ^ p < 0.1 \* p < 0.05, \*\* p < 0.01; \*\*\* p < 0.001

mPFC NR neural activity also depended on the size of reward delivered, which was significantly moderated by adolescent alcohol consumption (Figure 16D). We examined the relationship between mPFC’s NR population’s firing rate during the reward period, sex, alcohol, and number of sugar pellets delivered (Table 9, Model NRm1). Similar to Model PRm1, the number of sugar pellets delivered correlated with firing rate, but in the NR population this relationship was negative such that 1 sugar pellet decreased normalized firing at the time of reward by 0.31 compared to 0 pellets (b = −0.31, 95% CI [−0.36, −0.26], t (484) = −5.76, p < 0.001) and 3 sugar pellets decreased the firing rate by 0.46 compared to 0 pellets (b = −0.46, 95% CI [−0.51, −0.41], t (484) = −8.48, p < 0.001) and .15 compared to 1 pellet (b = −0.15, 95% CI [−0.20, −0.10], t (484) = −2.72, p = 0.007). There was no detectable relationship between firing rate during the reward period and sex (b = 0.04, 95% CI [−0.01, 0.09], t (484) = 0.70, p = 0.49) or alcohol (b = 0.02, 95% CI [0.00, 0.04], t (484) = 0.68, p = 0.50). In total, Model NRm1 explained 14% of the variance in firing rate (R^2^ = 0.14, F (4, 484) = 1.72, p < 0.001). Model NRm2 integrated the interaction between alcohol and sex into the model but it was not significant (b = −0.04, 95% CI [−0.12, 0.02], t (483) = −0.70, p = 0.49) nor was it a better fit than Model NRm1 (F (1, 483) = 0.48, p = 0.49). Model NPm3 examined the interaction between alcohol and number of sugar pellets. As the amount of alcohol consumed increased, there was a reduced decrease in the response to 3 pellet compared with 0 (b = 0.14, 95% CI [0.09, 0.19], t (482) = 2.84, p = 0.005). Thus the differences between large risky reward and omission observed in control animals diminishes to a lack of difference with increasing alcohol consumption. However, there was no difference between alcohol’s effect on 0 vs 1 pellet (b = 0.06, 95% CI [0.02, 0.12], t (482) = 1.34, p = 0.18) or 1 vs 3 pellets (b = 0.07, 95% CI [0.02, 0.12], t (482) = 1.50, p = 0.13). Model NRm3 explained 15% of the variance in firing rate during the reward period (R^2^ = 0.15, F (6, 482) = 14.09, p < 0.001) and was significantly better at explaining the variance than Model NRm1(F (2, 482) = 4.04, p = 0.02).

**Table 9.**
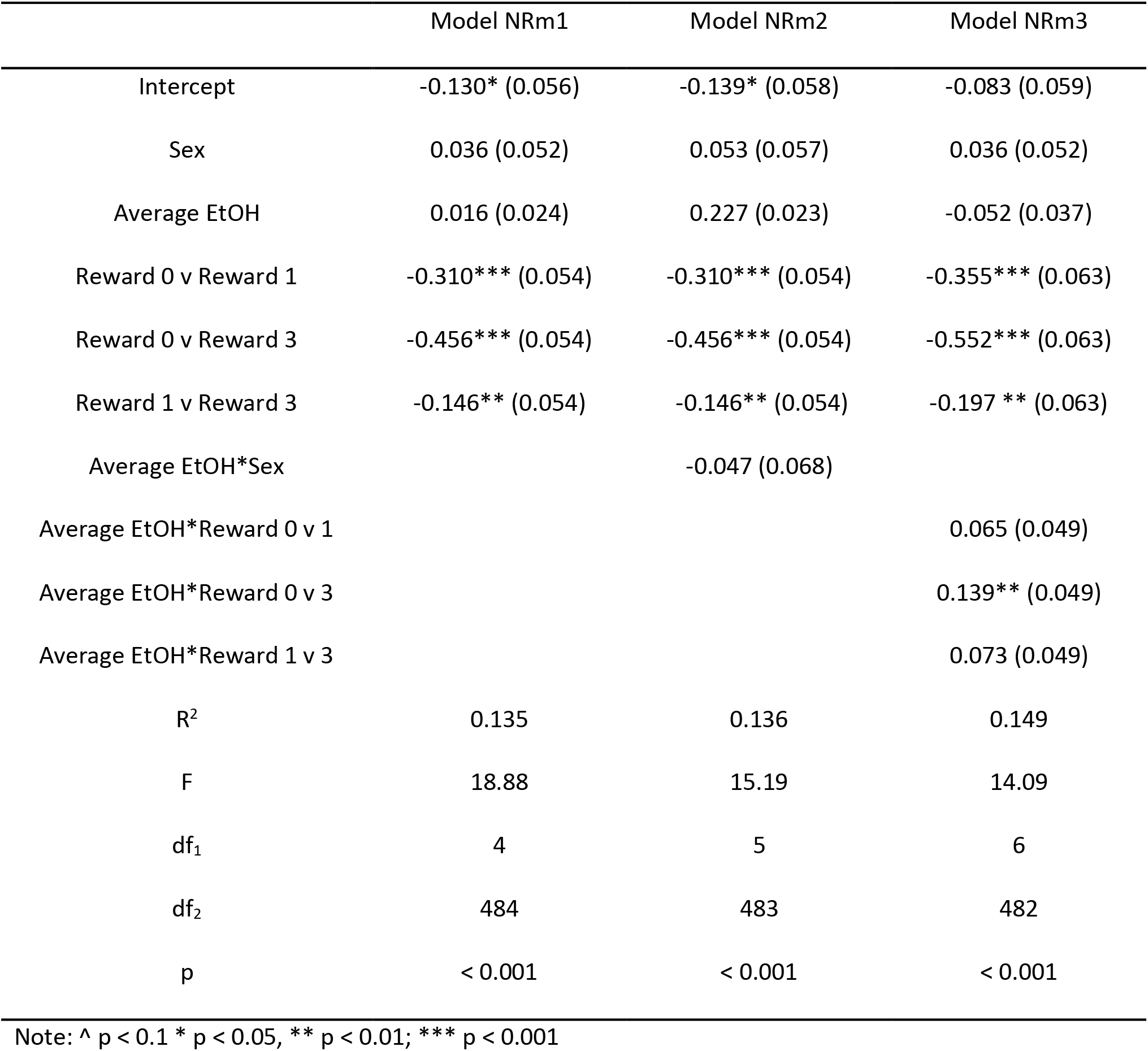
Summary of hierarchical regression analysis for variables predicting change in firing rate during the reward period in **mPFC NR** neurons.

Estrous cycle stage did not have an effect on baseline firing rate of mPFC PP and PR neurons (F (3, 100) = 0.94, p =0.42), nor did it effect mPFC NP and NR neurons’ baseline firing rate (F (3, 46) = 0.26, p = 0.85).

### Neural activity in OFC

Like mPFC, neural activity in OFC modulated at the time of the lever press and during reward delivery. Overall, OFC PP neurons showed a stereotyped increase at the time of lever press, regardless of which lever was selected and AIE (Figure 17A). The PP population of neurons in OFC were first examined for the relationships between firing rate at the time of lever press, sex, adolescent alcohol use, and lever pressed (Table 10, Model PPo1). This model explained 2% of the variance in firing rate at the time of lever press (R^2^ = 0.02, F (3, 274) = 1.39, p = 0.25) and neither sex (b = 0.11, 95% CI [−0.01, 0.23], t (274) = 0.79, p = 0.43), alcohol (b = −0.06, 95% CI [−0.12, 0.00], t (274) = −1.03, p = 0.30), or lever pressed (b = 0.14, 95% CI [0.02, 0.26], t (274) = 1.14, p = 0.26) had a main effect on firing rate. Model PPo2 tested the interaction between sex and adolescent alcohol use but the new term was not significant (b = 0.06, 95% CI [−0.10, 0.22], t (273) = 0.37, p = 0.71). The final model for OFC’s PP population, Model PPo3, probed if there was an interaction between alcohol and lever pressed on firing rate. However, this relationship did not appear to exist (b = 0.003, 95% CI [−0.11, 0.11], t (272) = 0.03, p = 0.98) and did not increase the model fit (R^2^ = 0.02, F (4, 273) = 1.04, p = 0.39).

**Figure 17.**
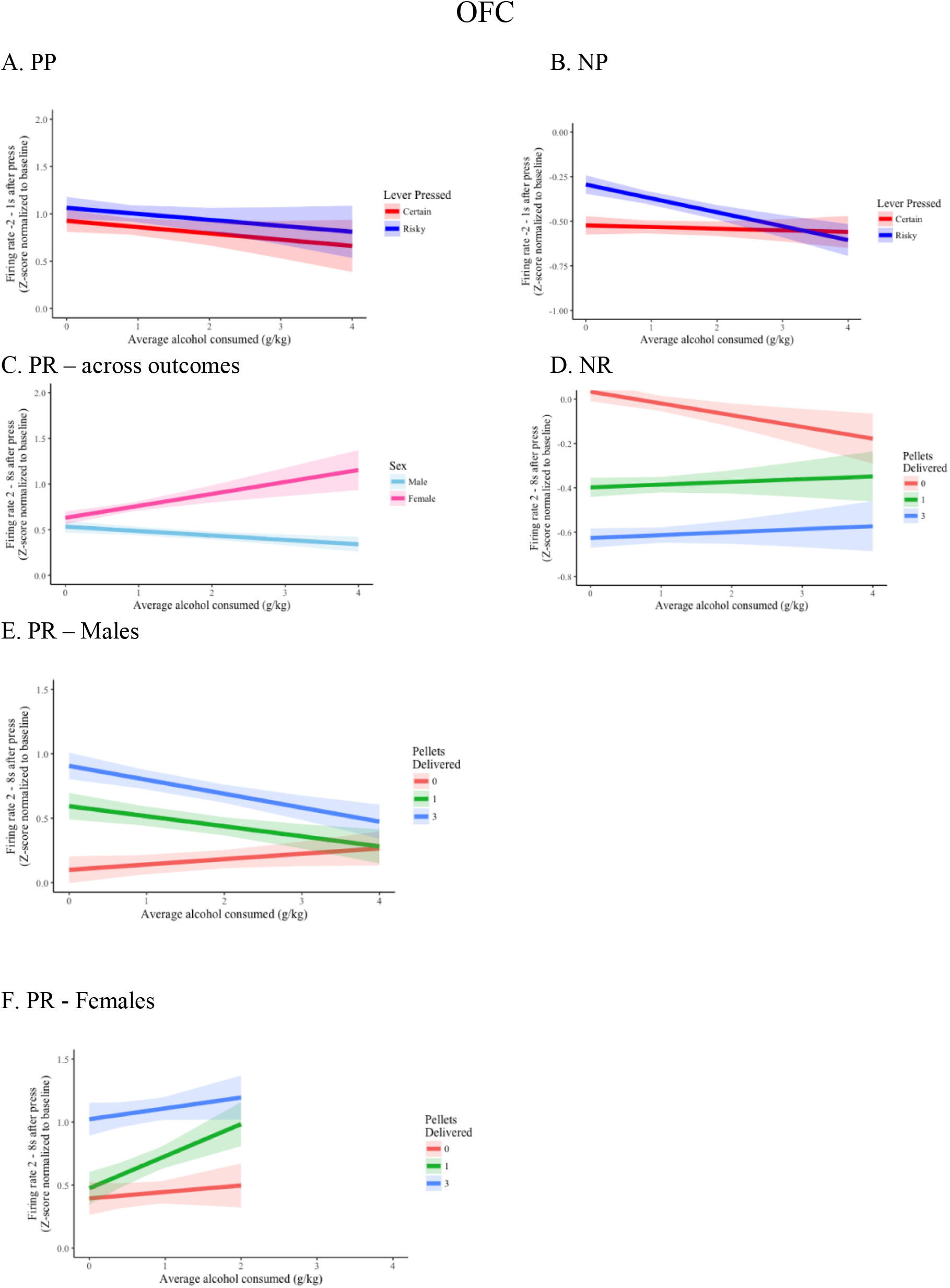
OFC responses. Simple slopes of chosen lever, moderated by alcohol consumed in adolescence, predicting firing rate at the time of lever press in PP population (A) and NP population (B). Simple slopes of sex, moderated by alcohol consumed in adolescence, predicting firing rate during the reward period in PR population (C). Simple slopes of reward size, moderated by alcohol consumed in adolescence, predicting firing rate during the reward period in NR population (D). Simple slopes of reward size, moderated by alcohol consumed in adolescence, predicting firing rate during the reward period in PR population in males (E) and females (F). Simple slopes of reward size, moderated by alcohol consumed in adolescence, predicting firing rate during the reward period in NR population (D).

**Table 10.** Summary of hierarchical regression analysis for variables predicting change in firing rate during lever press in **OFC PP** neurons.

|  | Model PPO1 | Model PPO2 | Model PPO3 |
| --- | --- | --- | --- |
| Intercept | 0.875*** (0.134) | 0.890*** (0.140) | 0.876*** (0.144) |
| Sex | 0.107 (0.136) | 0.071 (0.167) | 0.107* (0.136) |
| Average EtOH | -0.065 (0.063) | -0.076 (0.070) | -0.066 (0.085) |
| Lever Pressed | 0.140 (0.123) | 0.140 (0.122) | 0.137 (0.161) |
| Average EtOH*Sex |  | 0.061 (0.162) |  |
| Average EtOH*Lever Pressed |  |  | 0.003 (0.114) |
| R<sup>2</sup> | 0.015 | 0.015 | 0.015 |
| F | 1.389 | 1.295 | 1.038 |
| df<sub>1</sub> | 3 | 4 | 4 |
| df<sub>2</sub> | 274 | 273 | 273 |
| p | 0.247 | 0.370 | 0.388 |

Note: ^ p < 0.1 \* p < 0.05, \*\* p < 0.01; \*\*\* p < 0.001

The average responses of OFC NP neurons at the time of lever press are shown as a function of alcohol consumption in Figure 17B. We modeled the effects of sex, alcohol, and lever pressed on the firing rate during lever press in OFC’s NP neurons (Table 11, Model NPo1). Males had an higher firing rate during lever press than females (not shown; b = −0.12, 95% CI [−0.17, −0.07], t (204) = −2.10, p = 0.04), the risky lever caused an smaller decrease in firing compared to the certain lever (b = 0.14, 95% CI [0.09, 0.21], t (204) = 2.87, p = 0.005), and there was a trend toward increased alcohol causing a decrease in firing at the time of lever press (b = −0.04, 95% CI [−0.06, −0.02], t (204) = −1.94, p = 0.05). Overall the model explained 6% of the variance in firing rate at the time of lever press (R^2^ = 0.06, F (3, 204) = 4.69, p = 0.003). Model NPo2 tested for an interaction between sex and alcohol, but there was none (b = − 0.03, 95% CI [−0.08, 0.02], t (203) = −0.65, p = 0.52) and the model was not a better fit than Model NPo1 (F (1,203) = 0.42, p = 0.52). Model NPo3 tested if alcohol moderated the effect of lever pressed, resulting in a trend toward a significant, negative relationship (b = −0.07, 95% CI [−0.11, −0.03], t (203) = −1.69, p = 0.09). This suggests increased adolescent alcohol use decreases firing rate at the time of lever press more when the risky lever is pressed than the certain. However, this model obliterated the emerging main effect of alcohol (b = −0.009, 95% CI [−0.03, 0.03], t (203) = −0.31, p = 0.76) and was not significantly different than Model NPo1(F (1, 203) = 2.84, p = 0.09).

**Table 11.**
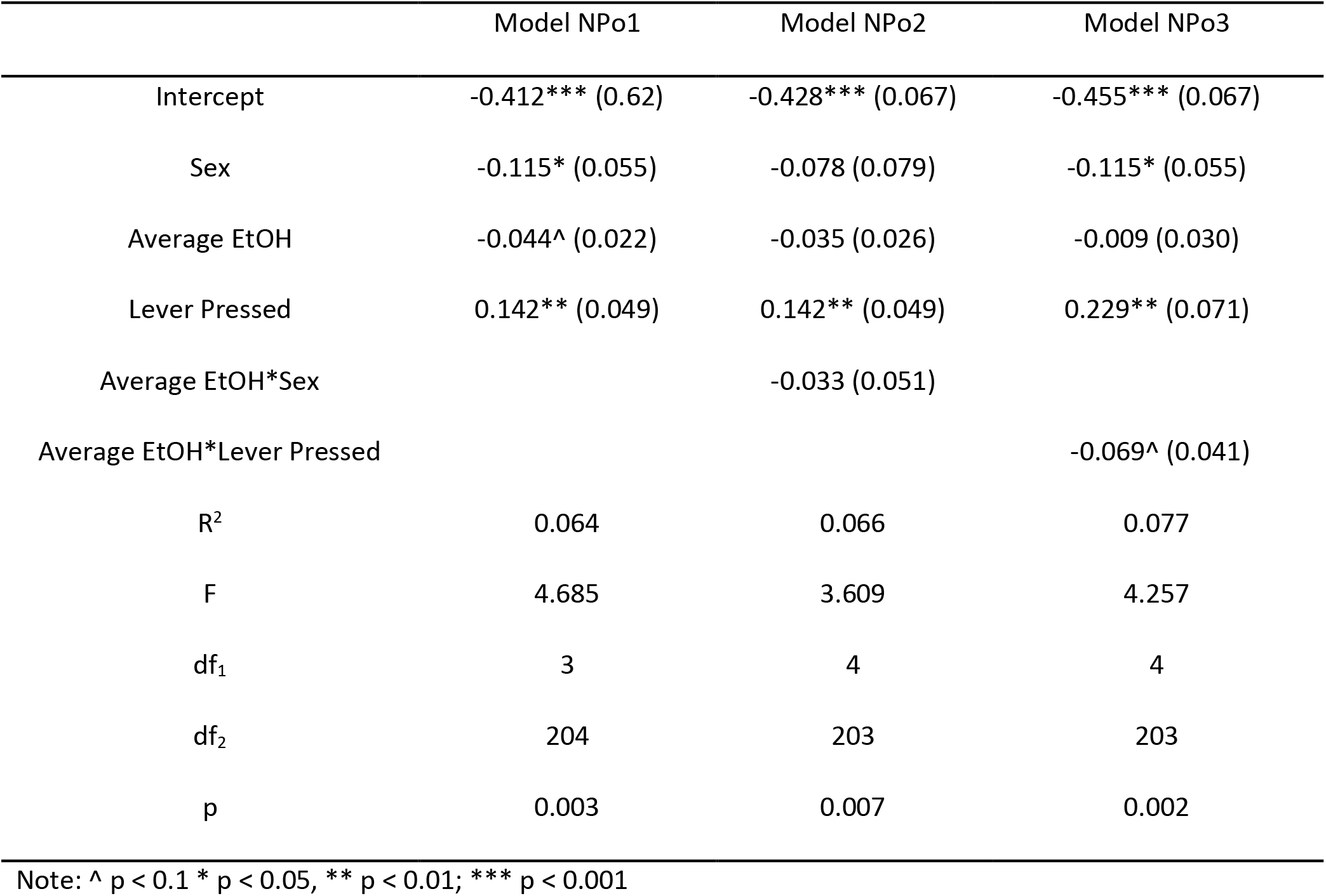
Summary of hierarchical regression analysis for variables predicting change in firing rate during lever press in **OFC NP** neurons.

OFC neurons that increased at the time of reward receipt, depended on reward size, sex, and alcohol consumption. The relationship between the activity of OFC’s PR population during reward delivery, sex, adolescent alcohol use, and number of sugar pellets delivered was modeled in Model PRo1 (Table 12). This model indicated that females had an increased firing rate compared to males (b = 0.26, 95% CI [0.19, 0.33], t (514) = 3.87, p < 0.001). There was a positive correlation between number of sugar pellets delivered and firing rate such that 1 sugar pellet increased firing by 0.27 compared to 0 pellets (b = 0.27, 95% CI [0.19, 0.33], t (514) = 3.87, p < 0.001), 3 sugar pellets increased firing by 0.61 compared to 0 pellets (b = 0.61, 95% CI [0.53, 0.69], t (514) = 7.80, p < 0.001), and 3 sugar pellets increased firing by 0.33 compared to 1 pellet (b = 0.33, 95% CI [0.25, 0.41], t (514) = 4.29, p < 0.001). Although, Model PRo1 did not reveal a main effect of alcohol (b = −0.02, 95% CI [−0.05, 0.01], t (514) = −0.66, p = 0.51), it did explain 14% of the variability in firing rate (R^2^ = 0.14, F (4, 514) = 20.08, p < 0.001). Model PRo2 revealed the interaction between sex and alcohol was significant (Figure 17C; b = 0.18, 95% CI [0.11, 0.25], t (513) = 2.46, p = 0.01) and was almost entirely responsible for the main effect of sex (b = 0.10, 95% CI [0.00, 0.20], t (513) = 2.46, p = 0.01). As alcohol consumption increased, neural activity during the reward period in adulthood increased significantly more in females compared to males. Additionally, Model PRo2 explained more of the variance in the firing rate than Model PRo1 (F (1, 513) = 6.06, p = 0.01).

**Table 12.** Summary of hierarchical regression analysis for variables predicting change in firing rate during the reward period in **OFC PR** neurons.

|  | Model PRo1 | Model PRo2 |
| --- | --- | --- |
| Intercept | 0.191* (0.076) | 0.240** (0.079) |
| Sex | 0.262*** (0.068) | 0.097 (0.095) |
| Average EtOH | -0.018 (0.027) | -0.048 (0.030) |
| Reward 0 v Reward 1 | 0.272*** (0.078) | 0.272*** (0.077) |
| Reward 0 v Reward 3 | 0.606*** (0.078) | 0.606*** (0.077) |
| Reward 1 v Reward 3 | 0.334*** (0.078) | 0.334*** (0.077) |
| Average EtOH*Sex |  | 0.179* (0.073) |
| R<sup>2</sup> | 0.135 | 0.145 |
| F | 20.08 | 17.43 |
| df<sub>1</sub> | 4 | 5 |
| df<sub>2</sub> | 514 | 513 |
| p | < 0.001 | < 0.001 |

Note: ^ p < 0.1 \* p < 0.05, \*\* p < 0.01; \*\*\* p < 0.001

The significant relationship between OFC PR responses with sex and alcohol consumption prompted the analysis of the sexes to be done separately for OFC’s PR population to avoid possible three-way interactions. Plots of average OFC PR neural responses to each reward size are shown for males (Figure 17E) and females (Figure 17F) separately. We re-evaluated the relationship between firing rate of OFC PR neurons in males, alcohol, and reward size (Table 13, Model PRo.M1). While the positive correlation between reward size and firing rate persisted (0 v 1: b = 0.30, 95% CI [0.20, 0.40], t (278) = 3.08, p = 0.002; 0 v 3: b = 0.56, 95% CI [0.46, 0.66], t (278) = 5.79, p < 0.001; 1 v 3: b = 0.26, 95% CI [0.16, 0.36], t (278) = 2.71, p = 0.01), a trend toward a negative effect of alcohol emerged as well (b = −0.05, 95% CI [− 0.08, −0.02], t (278) = −1.74, p = 0.08). Overall Model PRo.M1 was a significant predictor of firing rate during the reward period in male OFC PR neurons (R^2^ = 0.12, F (3, 278) = 12.20, p < 0.001). The interaction term between alcohol and reward size was incorporated into Model PRo.M2. This revealed that alcohol had a negative effect on firing rate when 3 pellets were delivered compared to 0 (b = −0.15, 95% CI [−0.22, −0.08], t (276) = −2.22, p = 0.03), and a trend toward a negative effect on firing rate when 1 pellet was delivered compared to 0 (b = −0.12, 95% CI [−0.19, −0.05], t (276) = −1.77, p = 0.08). Alcohol did not significantly moderate the effect of reward size on firing rate when 1 and 3 pellets were contrasted (b = −0.03, 95% CI [−0.10, 0.04], t (276) = −0.45, p = 0.66), indicating alcohol had a similar effect on the encoding of these rewards. Additionally, while Model PRo.M2 trended toward being a significantly better model than Model PRoM.1, it fell short in a statistical comparison (F (2, 276) = 2.75, p = 0.07). Analyses were repeated separately for female OFC PR neurons (Table 13, Model PRo.F1 and Model PRo.F2). In Model PRo.F1, when there was no interaction term, there was a trend toward a significant, positive effect of alcohol on firing rate (b = 0.13, 95% CI [0.06, 0.20], t (233) = 1.82, p = 0.07), and a positive contrast between 0 and 1 pellets’ effect on firing rate (b = 0.24, 95% CI [0.12, 0.36], t (233) = 1.90, p = 0.05). 3 sugar pellets also resulted in a greater increase in firing rate when compared to 0 (b = 0.66, 95% CI [0.54, 0.78], t (233) = 5.30, p < 0.001) or 1 pellet (b = 0.42, 95% CI [0.30, 0.54], t (233) = 3.37, p < 0.001). Cumulatively, Model PRo.F1 explained 12% of the variance in firing rate of the OFC PR population (R^2^ = 0.12, F (3, 233) = 10.69, p < 0.001). The incorporation of the interaction term into Model PRo.F2 did not result in any additional effects and was not a better model fit than Model PRo.F1 (F (2, 231) = 0.77, p = 0.47).

**Table 13.** Summary of hierarchical regression analysis for variables predicting change in firing rate during the reward period in **OFC PR** neurons.

|  | *Males* |  | *Females* |  |
| --- | --- | --- | --- | --- |
|  | Model PRo.M1 | Model PRo.M2 | Model PRo.F1 | Model PRo.F2 |
| Intercept | 0.244** (0.082) | 0.099 (0.103) | 0.331** (0.104) | 0.393** (0.131) |
| Average EtOH | -0.048^ (0.028) | 0.042 (0.048) | 0.1131^ (0.072) | 0.051 (0.124) |
| Reward 0 v Reward 1 | 0.300** (0.097) | 0.494*** (0.146) | 0.239* (0.124) | 0.080 (0.185) |
| Reward 0 v Reward 3 | 0.564*** (0.097) | 0.807*** (0.146) | 0.656*** (0.124) | 0.629*** (0.185) |
| Reward 1 v Reward 3 | 0.264*** (0.097) | 0.313* (0.146) | 0.417*** (0.124) | 0.548** (0.185) |
| Average EtOH* Reward 0 v 1 |  | -0.120^ (0.068) |  | 0.203 (0.176) |
| Average EtOH* Reward 0 v 3 |  | -0.150* (0.068) |  | 0.035 (0.176) |
| Average EtOH* Reward 1 v 3 |  | -0.030 (0.068) |  | -0.169 (0.176) |
| R<sup>2</sup> | 0.116 | 0.134 | 0.121 | 0.127 |
| F | 12.20 | 8.51 | 10.69 | 6.709 |
| df<sub>1</sub> | 3 | 5 | 3 | 5 |
| df<sub>2</sub> | 278 | 276 | 233 | 231 |
| p | < 0.001 | < 0.001 | < 0.001 | < 0.001 |

Note: ^ p < 0.1 \* p < 0.05, \*\* p < 0.01; \*\*\* p < 0.001

The subpopulation of NR neurons in OFC were modulated by reward size (Figure 17D). We examined sex, alcohol, and reward size as predictors of OFC’s NR population’s firing rate during the reward period (Table 14, Model NRo1). This revealed females had an overall lower firing rate during the reward period compared to males (not shown; b = −0.09, 95% CI [−0.13, −0.05], t (451) = −2.24, p = 0.03) and there was a negative relationship between reward size and firing rate (0 v 1: b = −0.38, 95% CI [−0.43, −0.33], t (451) = −7.89, p < 0.001; 0 v 3: b = −0.61, 95% CI [−0.66, −0.56], t (451) = −12.63, p < 0.001; 1 v 3: b = −0.23, 95% CI [−0.28, −0.18], t (451) = −4.74, p < 0.001). These factors explained 27% of the variability in firing rate (R^2^ = 0.27, F (4, 451) = 41.97, p < 0.001). Model NRo2 tested if alcohol moderated the effect of sex on firing rate but the interaction term was not significant (b = 0.02, 95% CI [−0.03, 0.07], t (450) = 0.44, p = 0.66) and did not pull any of the explained variability from the main effect of sex (b = −0.11, 95% CI [−0.16, −0.06], t (451) = −2.06, p = 0.04). Finally, Model NRo3 probed if alcohol moderated the effect of reward size. This did not result in any additional significant terms or an increase in explained variance compared to Model NRo1 (F (2, 449) = 1.32, p = 0.27).

**Table 14.** Summary of hierarchical regression analysis for variables predicting change in firing rate during the reward period in **OFC NR** neurons.

|  | Model NRo1 | Model NRo2 | Model NRo3 |
| --- | --- | --- | --- |
| Intercept | 0.049 (0.047) | 0.055 (0.049) | -0.084 (0.051) |
| Sex | -0.094* (0.042) | -0.107* (0.052) | -0.094* (0.042) |
| Average EtOH | -0.009 (0.020) | -0.015 (0.024) | -0.053 (0.034) |
| Reward 0 v Reward 1 | -0.379*** (0.048) | -0.379*** (0.048) | -0.431*** (0.061) |
| Reward 0 v Reward 3 | -0.607*** (0.048) | -0.607*** (0.048) | -0.660*** (0.061) |
| Reward 1 v Reward 3 | -0.228** (0.048) | -0.228** (0.048) | -0.229 ** (0.061) |
| Average EtOH*Sex |  | -0.020 (0.045) |  |
| Average EtOH*Reward 0 v 1 |  |  | 0.065 (0.047) |
| Average EtOH*Reward 0 v 3 |  |  | 0.065 (0.047) |
| Average EtOH*Reward 1 v 3 |  |  | 0.001 (0.047) |
| R<sup>2</sup> | 0.271 | 0.272 | 0.276 |
| F | 41.97 | 33.55 | 28.46 |
| df<sub>1</sub> | 4 | 5 | 6 |
| df<sub>2</sub> | 451 | 450 | 449 |
| p | < 0.001 | < 0.001 | < 0.001 |

Note: ^ p < 0.1 \* p < 0.05, \*\* p < 0.01; \*\*\* p < 0.001

Estrous cycle stage did not have an effect on baseline firing rate of OFC PP and PR neurons (F (3, 26) = 0.18, p =0.90), nor did it effect OFC NP and NR neurons’ baseline firing rate (F (3, 46) = 0.26, p = 0.85).

## DISCUSSION

We used an AIE exposure model to mimic voluntary moderate alcohol consumption in adolescence in order to look at the long-term effects on decision-making and neural encoding of decision variables. In our paradigm, control animals consumed approximately twice as much gelatin compared to alcohol animals (Figures 4 and 5). This is likely due to (1) the enhanced palatability of the control gelatin which was comprised of a greater polycose content, (2) a reduced preference for the experimental gelatin due to the bitter taste of alcohol, and (3) a limitation in consumption due to the intoxicating or sedative effects of alcohol. Based on prior work examining voluntary alcohol consumption (Bell et al., 2006; Maldonado, Finkbeiner, Alipour, & Kirstein, 2008; Vetter, Doremus-Fitzwater, & Spear, 2007) it is likely the former two justifications, as previous studies have successfully elicited alcohol consumption in excess to what was observed in this study. However, similar alcoholic gelatin models (McMurray et al., 2014; Peris et al., 2006) have observed alcohol intake comparable to what was documented in this study, reaffirming the method. There was a trend toward males consuming more gelatin than females, but this result wasn’t significant. Human data mirrors these findings such that there is no difference in the number of binge drinking episodes between adolescent boys and girls (Hamburg & Sharfstein, 2009; Schulte, Ramo, & Brown, 2009). Although daily consumption was normalized to weight to account for the discrepancy in size between sexes, this trend may indicate divergent patterns in escalation over time.

In the risk task, AIE had a negative, linear relationship with preference for the risky lever in both sexes (Figure 7). The effect of alcohol was most prominent in the 66% session, in which the risky lever was more greatly favored by low-alcohol consuming animals in this session than in the 16% and 33% (Figure S5). The incorporation of average alcohol use into regression models increased the significance of the variables sex and probability, indicating that some of the prior variability within these terms was actually attributable to alcohol use. Although sex was not a predictor of risk task performance in control animals, it was in alcohol animals and was consistent with the hypothesis that females are more risk averse than males. Furthermore, this phenotype appears to be amplified with alcohol as the slope of the regression line is steeper in females than males.

Sex, alcohol, and probability were all predicted to have an effect on decision-making behavior under uncertainty, but the directionality and magnitude of these variables’ impact was not congruent with our original expectations. The behavioral phenotypes observed in males in this study were discordant with results that have been replicated numerously (Boutros et al., 2015; Clark et al., 2012; McMurray et al., 2015; N. A. Nasrallah et al., 2011). Though the results were unexpected, differences in the task used here may account for these differences. Our risk task was a novel iteration of the traditional probabilistic discounting task, designed so animals were not over-trained and the sequence of probabilities was pseudorandomized. These are minor yet significant changes from previous studies which have trained rats on magnitude discrimination for up to 6 days (Clark et al., 2012; St Onge & Floresco, 2010) and the task for up to 24 days (St Onge & Floresco, 2010), until behavior was consistent. Furthermore, these experiments have utilized increasing or decreasing patterns of probabilistic discounting within a single session, despite evidence that that presentation order of probabilities can influence risky lever preference (Orsini et al., 2015). Thus, it is plausible that the unpredictable order of sessions combined with the lack of a well-practiced behavior strategy that has not been seen before to produce different behavioral results. In addition, we used a reward ratio (1 v 3 sugar pellets) that we have found to be more accurately discriminated (McMurray, Conway, & Roitman, 2017) compared to the ratio used in prior studies (McMurray et al., 2015). Performance on this task may instead more resemble a series of reversal learning tasks, which first introduces probabilistic rewards and then switches the location of their availability. In the face of contingencies that change multiple times, AIE animals may be more likely to adopt a strategy of choosing certain reward is calculation of probabilistic rewards is impaired. Lastly, one can speculate that the amount of alcohol used in prior reports, the mechanism of alcohol delivery, and the additional stressor of long-term social isolation of rats (which we did not introduce) could be confounding experimental factors. In conclusion, adolescent alcohol use does not universally cause increased risk preference in adulthood but does result in the inability to modulate behavior in response to changing contingencies.

While animals performed the risk task, we recorded from individual neurons in mPFC and OFC. Though these regions have been differentially thought to participate in the more immediate direction of behavior (mPFC) and the adaptation of behavior to changing reward contingencies (OFC), we observed neurons that responded both at the time of the behavioral choice (lever press) and during reward evaluation in both areas.

In addition to being engaged at the time of lever press, the PP population of mPFC appears to also be capable of discriminating choice-related cues. With this subpopulation, AIE consumption was related to a decrease in firing to the risky lever but not the certain (Figure 16A). This pattern suggests that the favored lever is signaled by greater changes in activity from baseline in this population of neurons, as the preference for the risky option decreased in animals with higher levels of adolescent alcohol consumption. The subpopulation of NP neurons in mPFC did not show any significant interaction between previous alcohol use and reduction in activity. This suggests that while mPFC’s PP and NP neuron populations are active at the time of lever press, they are encoding different properties of the decision-behavior.

The reward responsive neurons in mPFC similarly show a reduction in discrimination of reward size with increasing AIE. Both PR and NR neurons (Figure 16C & D) in mPFC differentiate between reward sizes in control animals, but with increasing adolescent alcohol intake, the responses to the three reward sizes become indiscriminable. This loss of differential reward encoding in mPFC corresponds with a bias toward the certain lever over either risky lever outcome, evocative of behavioral consequences of alcohol use seen in the risk task.

In OFC, none of the predicted factors had a discernible effect on the activity of the PP population in OFC at the time of lever press (Figure 17A), but the NP population did exhibit compelling patterns of activity. The negative impact of alcohol on firing rate appears to be mostly due to its negative moderation on the encoding of the risky lever (Figure 17B). This interaction is meaningful because it indicates that increased alcohol use correlates with a decrease in firing at the time of risky lever press, so that the normalized change in firing rate more closely resembles that of the certain lever.

OFC PR neurons were able to discriminate and encode reward size through graded increases in firing from baseline. However, adolescent alcohol use caused sexually dimorphic, long-lasting changes in this population’s neural activity (Figure 17C). In males, prior alcohol use was inversely related to the encoding of reward size (Figure 17E). This finding is similar to what was reported in McMurray et al., 2015 and supports the hypothesis that adolescent alcohol use disrupts OFC’s dynamic reward encoding that is essential for goal-directed behavior. In females, juvenile alcohol use resulted in an overall positive effect on the encoding of reward size, thus there was no differing interactions between alcohol and specific reward values. However, alcohol had the most substantial, positive effect on small certain rewards (Figure 17F). Consequentially, under control conditions, 1 sugar pellet was valued similarly to 0, but at higher levels of alcohol this relationship resulted in the encoding of 1 sugar pellet more closely resembling the encoding of 3 pellets. In summary, the sexually divergent data suggests that alcohol causes hypoactivity and the inability to encode economic value of rewards in males; however, adolescent alcohol use in females results in only fixed, definite rewards increasing in value. Both scenarios result in an overlap in the encoded value of larger and smaller rewards, yielding similar risk-averse phenotypes. In the NR population of neurons in OFC, reward size was negatively correlated with firing rate but did not show a dependence on alcohol consumption, suggesting this population is distinctly different from OFC’s PR population and might be protected from the toxic repercussions of alcohol.

The multitude of observed main effects and interactions modeled in the neural data speaks to the complexity of prefrontal cortex’s role in volition behavior and the extent of toxic assault it can endure.

## Notes

### Competing Interest Statement

The authors have declared no competing interest.

