## Supplemental Figures and Tables for "Effect of Voluntary Adolescent Alcohol Consumption on Encoding of Decision-Related Variables in Prefrontal Cortex"

**Supplemental Analyses**

A.

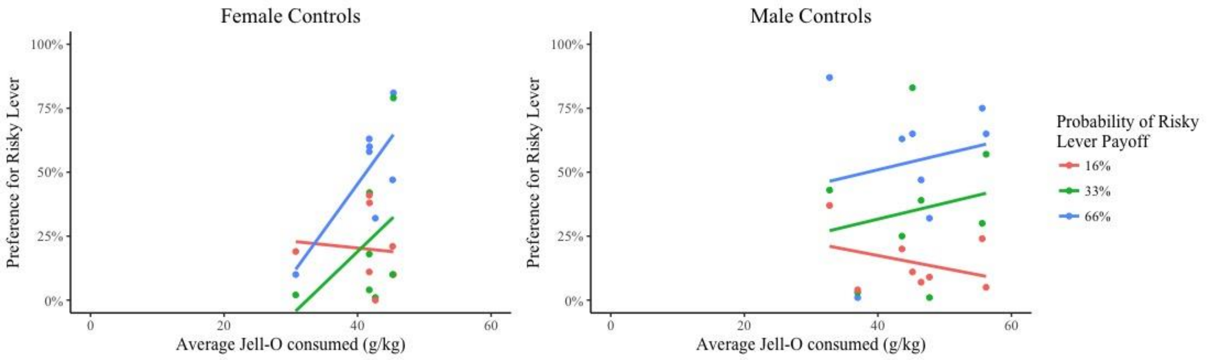


B.

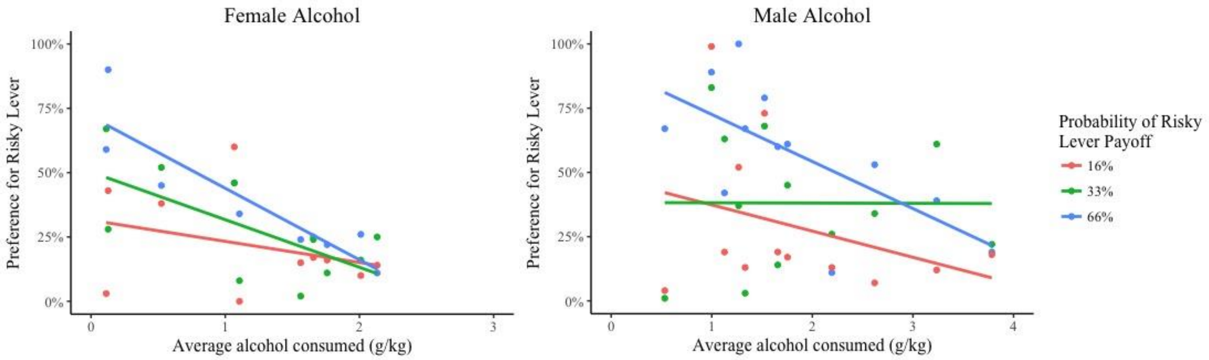


*Figure S1.* Simple slopes of the probability of risky lever payoff as a predictor of preference for the risky lever. (A) Control animals' performance on the risk task as predicted by probability of risky lever payoff and gelatin consumed in adolescence. (B) Alcohol animals' performance on the risk task as predicted by probability of risky lever payoff and prior alcohol use.

A.

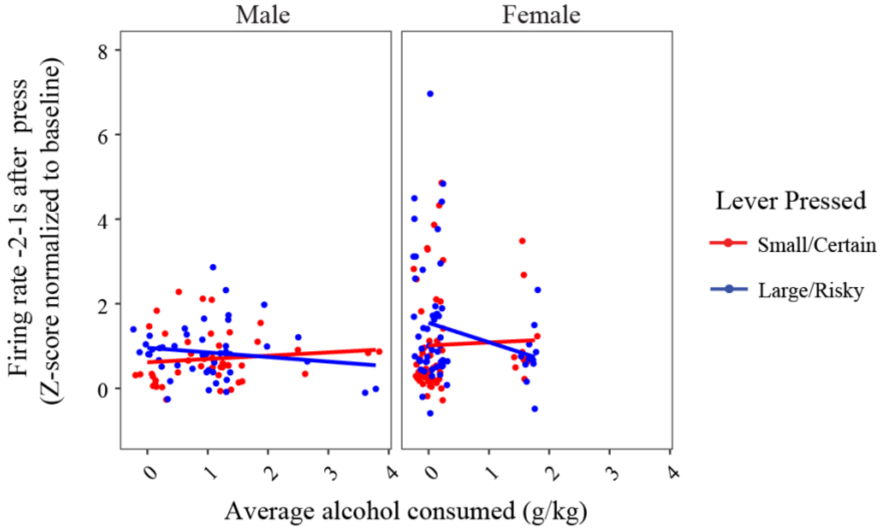


B.

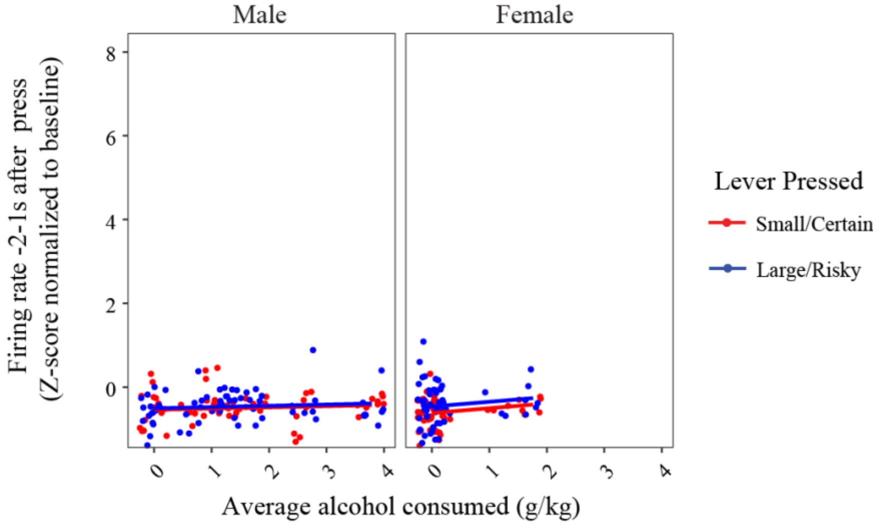


*Figure S2.* Patterns of neural activity in mPFC at the time of lever press as a factor of alcohol consumed in adolescence and lever pressed. (A) **PP neurons:** Change in normalized firing rate from baseline in neurons that on average exhibited an increase in firing rate at the time of lever press. (B) **NP neurons:** Change in normalized firing rate from baseline in neurons that on average exhibited a decrease in firing rate at the time of lever press.

Table S1

*Summary of mixed linear regression analysis for variables predicting change in firing rate during lever press in* ***mPFC PP*** *neurons.*

| | Model PPm1 | Model PPm2 | Model PPm3 |
|---|---|---|---|
| **Fixed Effects** | | | |
| Intercept | 0.723*** (0.177) | 0.648*** (0.199) | 0.623*** (0.182) |
| Sex | 0.386* (0.182) | 0.494* (0.225) | 0.386* (0.182) |
| Average EtOH | -0.090 (0.109) | -0.016 (0.142) | 0.072 (0.130) |
| Lever Pressed | 0.276* (0.120) | 0.276* (0.120) | 0.476** (0.146) |
| Average EtOH*Sex | | -0.179 (0.221) | |
| Average EtOH*Lever Pressed | | | -0.325* (0.141) |
| **Random Effect** | | | |
| N group | 108 | 108 | 108 |
| Observations | 216 | 216 | 216 |

Note: ^ $p < 0.1$ * $p < 0.05$, ** $p < 0.01$; *** $p < 0.001$

Table S2

*Summary of mixed linear regression analysis for variables predicting change in firing rate during lever press in **mPFC NP** neurons.*

| | Model NPm1 | Model NPm2 | Model NPm3 |
|---|---|---|---|
| **Fixed Effects** | | | |
| Intercept | -0.586*** (0.056) | -0.568*** (0.058) | -0.597*** (0.059) |
| Sex | 0.018 (0.060) | -0.017 (0.067) | 0.018 (0.060) |
| Average EtOH | 0.042 (0.026) | 0.030 (0.028) | 0.055 (0.033) |
| Lever Pressed | 0.092^ (0.047) | 0.092^ (0.047) | 0.115^ (0.060) |
| Average EtOH*Sex | | 0.084 (0.074) | |
| Average EtOH*Lever Pressed | | | -0.026 (0.041) |
| **Random Effect** | | | |
| N group | 126 | 126 | 126 |
| Observations | 252 | 252 | 252 |

Note: ^ $p < 0.1$ * $p < 0.05$, ** $p < 0.01$; *** $p < 0.001$

A.

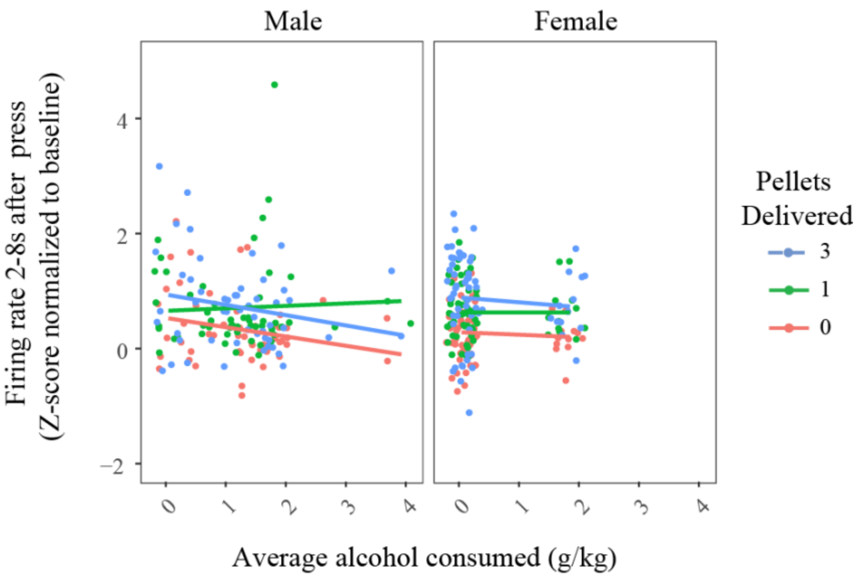


B.

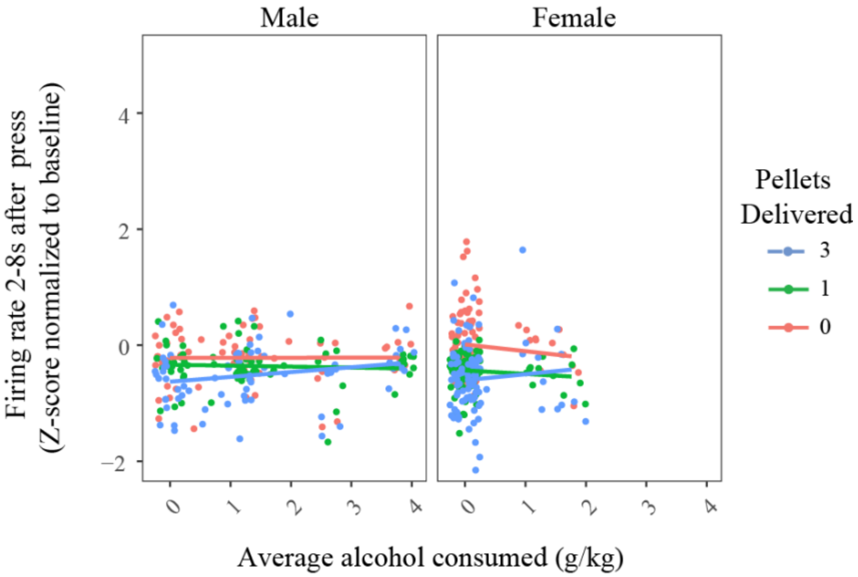


*Figure S3.* Patterns of neural activity in mPFC during the reward period as a factor of alcohol consumed in adolescence and lever pressed. (A) **PR neurons:** Change in normalized firing rate from baseline in neurons that on average exhibited an increase in firing rate during the reward period. (B) **NR neurons:** Change in normalized firing rate from baseline in neurons that on average exhibited a decrease in firing rate during the reward period.

Table S3

*Summary of mixed linear regression analysis for variables predicting change in firing rate during the reward period in* ***mPFC PR*** *neurons.*

| | Model PRm1 | Model PRm2 | Model PRm3 |
|---|---|---|---|
| **Fixed Effects** | | | |
| Intercept | 0.388*** (0.099) | 0.417*** (0.114) | 0.397*** (0.105) |
| Sex | -0.066 (0.095) | -0.107 (0.124) | -0.066 (0.095) |
| Average EtOH | -0.076 (0.056) | -0.102 (0.074) | -0.088 (0.074) |
| Reward 0 v Reward 1 | 0.366*** (0.073) | 0.366*** (0.073) | 0.286** (0.095) |
| Reward 0 v Reward 3 | 0.496*** (0.073) | 0.496*** (0.073) | 0.550*** (0.095) |
| Reward 1 v Reward 3 | 0.130^ (0.073) | 0.130^ (0.073) | 0.264** (0.095) |
| Average EtOH*Sex | | 0.058 (0.112) | |
| Average EtOH*Reward 0 v 1 | | | 0.111 (0.085) |
| Average EtOH*Reward 0 v 3 | | | -0.076 (0.085) |
| Average EtOH*Reward 1 v 3 | | | -0.187* (0.085) |
| **Random Effects** | | | |
| N group | 115 | 115 | 115 |
| Observations | 345 | 345 | 345 |

Note: ^ p < 0.1 * p < 0.05, ** p < 0.01; *** p < 0.001

Table S4

*Summary of mixed linear regression analysis for variables predicting change in firing rate during the reward period in* ***mPFC NR*** *neurons.*

| | Model NRm1 | Model NRm2 | Model NRm3 |
|---|---|---|---|
| **Fixed Effects** | | | |
| Intercept | -0.130 * (0.060) | -0.139* (0.062) | -0.083 (0.063) |
| Sex | 0.036 (0.060) | -0.053 (0.066) | 0.036 (0.061) |
| Average EtOH | 0.016 (0.027) | -0.023 (0.029) | -0.052 (0.037) |
| Reward 0 v Reward 1 | -0.310*** (0.049) | -0.310*** (0.049) | -0.355** (0.057) |
| Reward 0 v Reward 3 | -0.456*** (0.049) | -0.456*** (0.049) | -0.552*** (0.057) |
| Reward 1 v Reward 3 | -0.146** (0.049) | -0.146**(0.049) | -0.197***(0.057) |
| Average EtOH*Sex | | -0.047 (0.078) | |
| Average EtOH*Reward 0 v 1 | | | 0.065 (0.044) |
| Average EtOH*Reward 0 v 3 | | | 0.139** (0.044) |
| Average EtOH*Reward 1 v 3 | | | 0.073^ (0.044) |
| **Random Effects** | | | |
| N group | 163 | 163 | 163 |
| Observations | 489 | 489 | 489 |

Note: ^ $p < 0.1$ * $p < 0.05$, ** $p < 0.01$; *** $p < 0.001$

A.

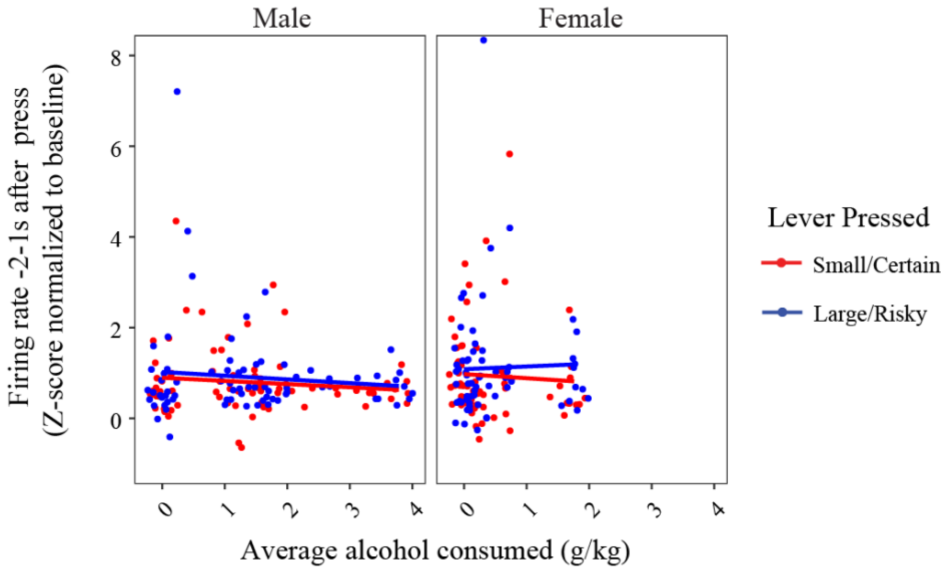


B.

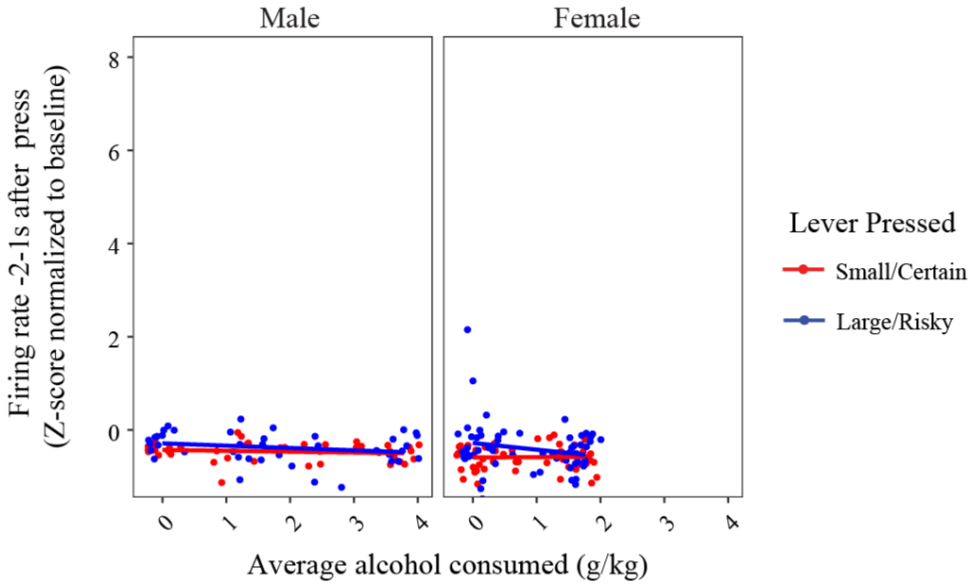


*Figure S4.* Patterns of neural activity in OPFC at the time of lever press as a factor of alcohol consumed in adolescence and lever pressed. (A) **PP neurons:** Change in normalized firing rate from baseline in neurons that on average exhibited an increase in firing rate at the time of lever press. (B) **NP neurons:** Change in normalized firing rate from baseline in neurons that on average exhibited a decrease in firing rate at the time of lever press.

Table S5

*Summary of mixed linear regression analysis for variables predicting change in firing rate during lever press in* ***OFC PP*** *neurons.*

| | Model PPo1 | Model PPo2 | Model PPo3 |
|---|---|---|---|
| **Fixed Effects** | | | |
| Intercept | 0.875*** (0.159) | 0.890*** (0.167) | 0.876*** (0.161) |
| Sex | 0.107 (0.178) | 0.071 (0.217) | 0.106 (0.178) |
| Average EtOH | -0.065 (0.082) | -0.076 (0.091) | -0.066 (0.087) |
| Lever Pressed | 0.140* (0.063) | 0.140* (0.063) | 0.137^ (0.082) |
| Average EtOH*Sex | | 0.060 (0.211) | |
| Average EtOH*Lever Pressed | | | 0.003 (0.058) |
| **Random Effect** | | | |
| N group | 139 | 139 | 139 |
| Observations | 278 | 278 | 278 |

Note: ^ $p < 0.1$ * $p < 0.05$, ** $p < 0.01$; *** $p < 0.001$

Table S6

*Summary of mixed linear regression analysis for variables predicting change in firing rate during lever press in* ***OFC NP*** *neurons.*

| | Model NPo1 | Model NPo2 | Model NPo3 |
|---|---|---|---|
| **Fixed Effects** | | | |
| Intercept | -0.412*** (0.063) | -0.428*** (0.068) | -0.455*** (0.067) |
| Sex | -0.115* (0.056) | -0.078 (0.081) | -0.115* (0.056) |
| Average EtOH | -0.044^ (0.023) | -0.035 (0.027) | -0.009 (0.030) |
| Lever Pressed | 0.142** (0.047) | 0.142** (0.047) | 0.229** (0.067) |
| Average EtOH*Sex | | -0.033 (0.052) | |
| Average EtOH*Lever Pressed | | | -0.069^ (0.038) |
| **Random Effect** | | | |
| N group | 104 | 104 | 104 |
| Observations | 208 | 208 | 208 |

Note: ^ p < 0.1 * p < 0.05, ** p < 0.01; *** p < 0.001

A.

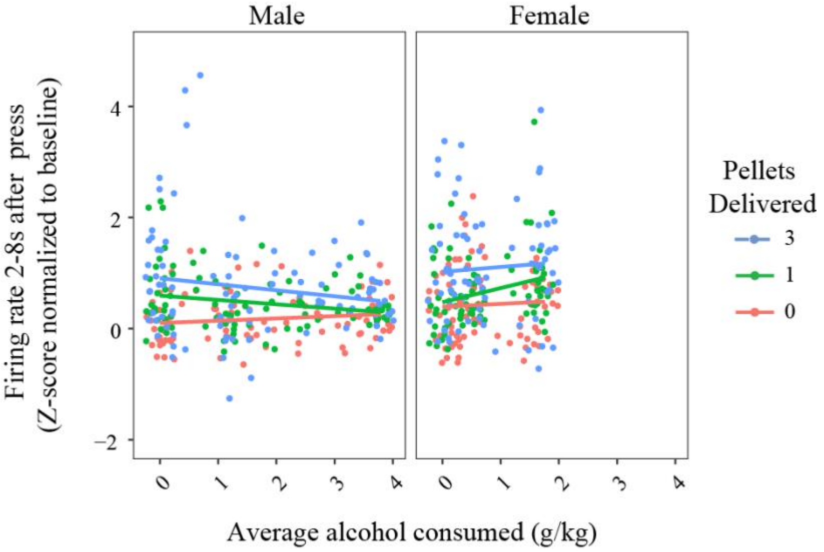


B.

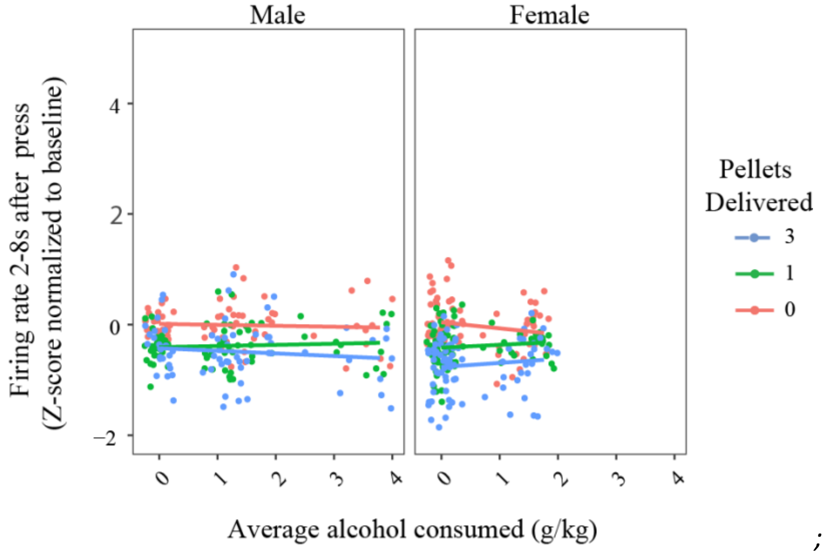


;

*Figure S5.* Patterns of neural activity in OFC during the reward period as a factor of alcohol consumed in adolescence and lever pressed. (A) **PR neurons:** Change in normalized firing rate from baseline in neurons that on average exhibited an increase in firing rate during the reward period. (B) **NR neruons:** Change in normalized firing rate from baseline in neurons that on average exhibited a decrease in firing rate during the reward period.

Table S7

*Summary of mixed linear regression analysis for variables predicting change in firing rate during the reward period in **OFC PR** neurons.*

| | Model PRo1 | Model PRo2 |
|---|---|---|
| **Fixed Effects** | | |
| Intercept | 0.191* (0.085) | 0.240** (0.088) |
| Sex | 0.262** (0.083) | 0.097 (0.116) |
| Average EtOH | -0.018 (0.034) | -0.048 (0.036) |
| Reward 0 v Reward 1 | 0.272*** (0.067) | 0.272*** (0.067) |
| Reward 0 v Reward 3 | 0.606*** (0.067) | 0.606*** (0.067) |
| Reward 1 v Reward 3 | 0.334*** (0.067) | 0.334*** (0.067) |
| Average EtOH*Sex | | 0.179* (0.089) |
| **Random Effect** | | |
| N group | 173 | 173 |
| Observations | 519 | 519 |

Note: ^ p < 0.1 * p < 0.05, ** p < 0.01; *** p < 0.001

Table S8

*Summary of mixed linear regression analysis for variables predicting change in firing rate during the reward period in **OFC PR** neurons continued.*

| | *Males* | | *Females* | |
|---|---|---|---|---|
| | Model PRo.M1 | Model PRo.M2 | Model PRo.F1 | Model PRo.F2 |
| **Fixed Effects** | | | | |
| Intercept | 0.244** (0.087) | 0.099 (0.102) | 0.331** (0.111) | 0.393** (0.129) |
| Average EtOH | -0.048 (0.033) | 0.042 (0.047) | 0.131 (0.089) | 0.051 (0.123) |
| Reward 0 v Reward 1 | 0.300*** (0.085) | 0.494*** (0.126) | 0.239* (0.104) | 0.080 (0.155) |
| Reward 0 v Reward 3 | 0.564*** (0.085) | 0.807*** (0.126) | 0.656*** (0.104) | 0.629*** (0.155) |
| Reward 1 v Reward 3 | 0.264*** (0.085) | 0.313*(0.126) | 0.417 *** (0.104) | 0.548*** (0.155) |
| Average EtOH* Reward 0 v 1 | | -0.120* (0.058) | | 0.203 (0.146) |
| Average EtOH* Reward 0 v 3 | | -0.150* (0.058) | | 0.035 (0.147) |
| Average EtOH* Reward 1 v 3 | | -0.030 (0.058) | | -0.169 (0.147) |
| **Random Effect** | | | | |
| N group | 94 | 94 | 79 | 79 |
| Observations | 282 | 282 | 273 | 273 |

Note: ^ $p < 0.1$ * $p < 0.05$, ** $p < 0.01$; *** $p < 0.001$

Table S9

*Summary of hierarchical regression analysis for variables predicting change in firing rate during the reward period in* ***OFC NR*** *neurons.*

| | Model NRo1 | Model NRo2 | Model NRo3 |
|---|---|---|---|
| **Fixed Effects** | | | |
| Intercept | 0.049 (0.051) | 0.055 (0.054) | -0.084 (0.054) |
| Sex | -0.094^ (0.049) | -0.107^ (0.061) | -0.094^ (0.049) |
| Average EtOH | -0.009 (0.024) | -0.015 (0.028) | -0.053 (0.034) |
| Reward 0 v Reward 1 | -0.379*** (0.043) | -0.379*** (0.043) | -0.431*** (0.054) |
| Reward 0 v Reward 3 | -0.607*** (0.043) | -0.607*** (0.043) | -0.660*** (0.054) |
| Reward 1 v Reward 3 | -0.228** (0.043) | -0.228** (0.043) | -0.229 *** (0.054) |
| Average EtOH*Sex | | -0.020 (0.053) | |
| Average EtOH*Reward 0 v 1 | | | 0.065 (0.041) |
| Average EtOH*Reward 0 v 3 | | | 0.066 (0.041) |
| Average EtOH*Reward 1 v 3 | | | 0.001 (0.041) |
| **Random Effect** | | | |
| N group | 152 | 152 | 152 |
| Observations | 456 | 456 | 456 |

Note: ^ $p < 0.1$ * $p < 0.05$, ** $p < 0.01$; *** $p < 0.001$
